# Multi-color single molecule imaging uncovers extensive heterogeneity in mRNA decoding

**DOI:** 10.1101/477661

**Authors:** Sanne Boersma, Deepak Khuperkar, Bram M.P. Verhagen, Stijn Sonneveld, Jonathan B. Grimm, Luke D. Lavis, Marvin E. Tanenbaum

## Abstract

mRNA translation is a key step in decoding genetic information. Genetic decoding is surprisingly heterogeneous, as multiple distinct polypeptides can be synthesized from a single mRNA sequence. To study translational heterogeneity, we developed the MoonTag, a new fluorescence labeling system to visualize translation of single mRNAs. When combined with the orthogonal SunTag system, the MoonTag enables dual readouts of translation, greatly expanding the possibilities to interrogate complex translational heterogeneity. By placing MoonTag and SunTag sequences in different translation reading frames, each driven by distinct translation start sites, start site selection of individual ribosomes can be visualized in real-time. We find that start site selection is largely stochastic, but that the probability of using a particular start site differs among mRNA molecules, and can be dynamically regulated over time. Together, this study provides key insights into translation start site selection heterogeneity, and provides a powerful toolbox to visualize complex translation dynamics.

## Introduction

Translation of mRNAs by ribosomes is a key step in decoding the genetic information stored in DNA and mRNA, and regulation of translation plays an important role in shaping the proteome (Hinnebusch et al., 2016; Jackson et al., 2010; Schwanhäusser et al., 2009). Typically, translation initiates at the most upstream (i.e. most 5’) translation start codon, usually an AUG codon, and then continues in the same reading frame until it encounters the first in-frame stop codon (here referred to as canonical translation). However, more recent work has shown that translation of many, if not most mRNAs, is far more complex, and that different regions of an mRNA can be translated. For example, many mRNAs contain multiple open reading frames, including upstream ORFs (uORFs), which are short ORFs upstream of the ‘main’ ORF that generally repress translation of the main ORF (Calvo et al., 2009; Chew et al., 2016; Johnstone et al., 2016; Wethmar, 2014). Moreover, ribosomes can translate each nucleotide sequence in 3 different reading frames, resulting in 3 completely unrelated polypeptides (Atkins et al., 2016). Ribosomes translating some eukaryotic or viral RNAs can also undergo frameshifting, changing the reading frame during translation elongation (Dinman, 2012; Dunkle and Dunham, 2015). Finally, ribosomes can bypass stop codons under certain conditions to generate C-terminally extended proteins (Beznosková et al., 2015; Dunn et al., 2013; Schueren and Thoms, 2016). While many examples are known where non-canonical translation occurs productively to generate functional proteome diversity, it is important to note that non-canonical translation may also occur inappropriately, due to errors in translation. Such errors likely result in synthesis of misfolded and/or dysfunctional polypeptides, which may inhibit the function of the natively folded protein, and can cause proteotoxic stress to the cell.

Critical for determining the translated region of the mRNA, is the selection of the correct translation start site. In eukaryotes, the translation start site is selected during a process in which the 43S translation pre-initiation complex, including the small ribosomal subunit, ‘scans’ along the mRNA in a 5’-to-3’ direction in search of a start codon (Aitken and Lorsch, 2012; Hinnebusch et al., 2016; Jackson et al., 2010). The identification of the correct start site for a scanning ribosome is complex, as; 1) many genes contain one or more AUG sequences in their 5’ untranslated region (UTR) (Iacono et al., 2005); 2) translation can also initiate, albeit generally less efficiently, on near-cognate start codons (e.g. GUG, CUG) (Gao et al., 2015; Ingolia et al., 2011; Kearse and Wilusz, 2017; Lee et al., 2012; Na et al., 2018); 3) the canonical start site may not be recognized with 100% efficiency (Kozak, 1986; Lind and Aqvist, 2016; Pisareva and Pisarev, 2016); 4) after translating of a short ORF (e.g. a uORF), a ribosome can reinitiate translation at a downstream start site, thus initiating at multiple start sites on a single mRNA molecule (Barbosa et al., 2013; Hinnebusch et al., 2016; Somers et al., 2013; Starck et al., 2016; Wethmar, 2014). An additional layer of complexity in selection of a start site is the existence of multiple different transcript isoforms for many genes. For example, alternative transcription start site (TSS) usage or alternative splicing could create different transcript isoforms, and these isoforms may contain a different set of translation start sites (Floor and Doudna, 2016; Wang et al., 2016b).

While ensemble measurements have identified multiple translation start sites for many genes, it is currently unclear whether all start sites are used on each individual mRNA molecules, and if so, how their relative usage is regulated. In the simplest model, ribosomes initiate translation on each possible start site with a pre-defined probability, which depends on the sequence of the start codon and its local sequence context (i.e. Kozak consensus sequence). In this model, all possible start sites are used on each mRNA molecule and translation start site selection by the scanning pre-initiation complex is purely stochastic. Alternatively, relative start site usage could vary among different mRNA copies originating from the same gene, for example due to differences in the transcript isoforms, RNA structure or due to regulatory factors, such as RNA binding proteins (RBPs) or RNA modifications. Regulation of start site usage would provide an intriguing possibility to tune protein levels as well as protein sequence in space and time. Recently, the m6A mRNA modification was shown to bias translation start site selection (Zhou et al., 2018), indicating that regulation of translation start site selection may occur, at least for a subset of transcripts.

While the mechanisms of canonical translation have been extensively studied, the prevalence and underlying causes of heterogeneity in mRNA translation have remained largely unexplored. Currently used technologies, like ribosome profiling and GFP or luciferase reporters, are not ideally suited to detect variability of mRNA translation, including variability in translation start site selection, because 1) they cannot distinguish which translation start sites are used on which mRNA molecules, or whether multiple start sites are used on individual mRNA molecules; 2) they cannot track translation start site usage in space and time for individual mRNA molecules; 3) it is challenging to detect infrequently used start sites above the experimental noise, especially if many different infrequently used start sites exist in an mRNA; 4) static measurements may not readily detect start sites that trigger mRNA degradation. Start sites that result in out-of-frame translation, which likely represent the majority of non-canonical translation initiation events, may trigger nonsense mediated-mRNA decay (Lykke-Andersen and Jensen, 2015), resulting in rapid decay of the mRNAs that preferentially use such alternative start sites. Therefore, new tools are required to provide real-time readouts of translational heterogeneity and translation start site selection on single mRNAs.

We have recently developed a fluorescence labeling strategy, called SunTag, consisting of a genetically-encoded fluorescently-labeled intracellular antibody and a peptide epitope (Tanenbaum et al., 2014). We and others have shown that the SunTag system (Wang et al., 2016a; Wu et al., 2016; Yan et al., 2016), or a labeling system with a purified antibody (Morisaki et al., 2016), can be applied to fluorescently label nascent polypeptides, enabling visualization of translation of individual mRNA molecules over time. However, the SunTag system only provides a single read-out of translation and is therefore not suited to study more complex translation events. Here, we develop an orthogonal genetically-encoded system for labeling nascent polypeptides, which we call the MoonTag. We combine the MoonTag and SunTag systems in a single mRNA, allowing dual-color readouts of translation, which provides an opportunity to study non-canonical translation and translational heterogeneity. We show that a reporter mRNA containing SunTag and MoonTag sequences in different translation reading frames, can report on translation start site selection of individual ribosomes in real-time. Using this system, we study translational heterogeneity on a variety of different types of mRNAs, including mRNAs containing a uORF. We find that 1) multiple translation start sites are used intermittently on most mRNAs, 2) the usage frequency of these translation start sites differs substantially among mRNA molecules originating from the same gene and 3) translation start site usage can be dynamically regulated over time on single mRNAs. Taken together, our study reveals translation start site selection dynamics and heterogeneity, and provides a generally applicable method for studying complex translation dynamics.

## Results

### Development of the MoonTag, a new fluorescence labeling system to visualize translation of single mRNAs

To obtain multiple readouts of translation of single mRNA molecules in real-time, we aimed to establish a second genetically-encoded antibody-epitope pair for nascent chain labeling (Fig. 1A), orthogonal to our previously developed SunTag system (Tanenbaum et al., 2014). Previously, antibodies targeting the HA-and Flag-tags have been developed for translation imaging (Morisaki et al., 2016; Viswanathan et al., 2015). However, these systems require introduction of purified, *in vitro* labeled antibody fab fragments into cells, and have not been genetically encoded for stable expression in cells. To identify potential antibody-peptide pairs that could be genetically encoded and used for *in vivo* nascent chain labeling, we performed a literature search for single chain antibodies (e.g. nanobodies) that bound a linear epitope with high affinity, and identified seven antibody-peptides pairs (See Methods for details). To test whether antibody-peptide binding was retained inside living cells, we fused the antibodies to GFP, and fused each peptide in twelve tandem copies to mCherry. We tethered the 12xpeptide-mCherry fusions to the outer surface of the mitochondria through fusion to the mitochondrial targeting domain (referred to as Mito) of the outer mitochondrial membrane protein MitoNEET, as described previously (Tanenbaum et al., 2014). If an antibody binds to its cognate epitope in cells, GFP and mCherry signals are expected to co-localize at the outer surface of mitochondria. In all seven cases, the Mito-mCherry-12xpeptide fusions localized to the mitochondria and the antibody-GFP fusions were expressed in the cytoplasm. However, a clear enrichment of GFP signal near mitochondria was observed for only a single antibody-peptide pair (Fig. 1B, gp41 nanobody (Nb-gp41-GFP)). Nb-gp41-GFP was localized diffusely in the cytoplasm in the absence of the peptide (Fig. 1C), confirming that GFP enrichment near mitochondria reflects antibody-peptide binding. The gp41 peptide is a 15 amino acid peptide from the HIV envelope protein complex subunit gp41. In the peptide array the gp41 peptide repeats are separated by 11 amino acid glycine-serine linkers. The gp41 antibody is a 123 amino acid Llama VHH nanobody (clone 2H10), which binds the peptide *in vitro* with an affinity of ~30nM (Lutje Hulsik et al., 2013). As this antibody-peptide system is orthogonal to our ‘SunTag’ system, we refer to it as the ‘MoonTag’ system.

**Figure 1.**
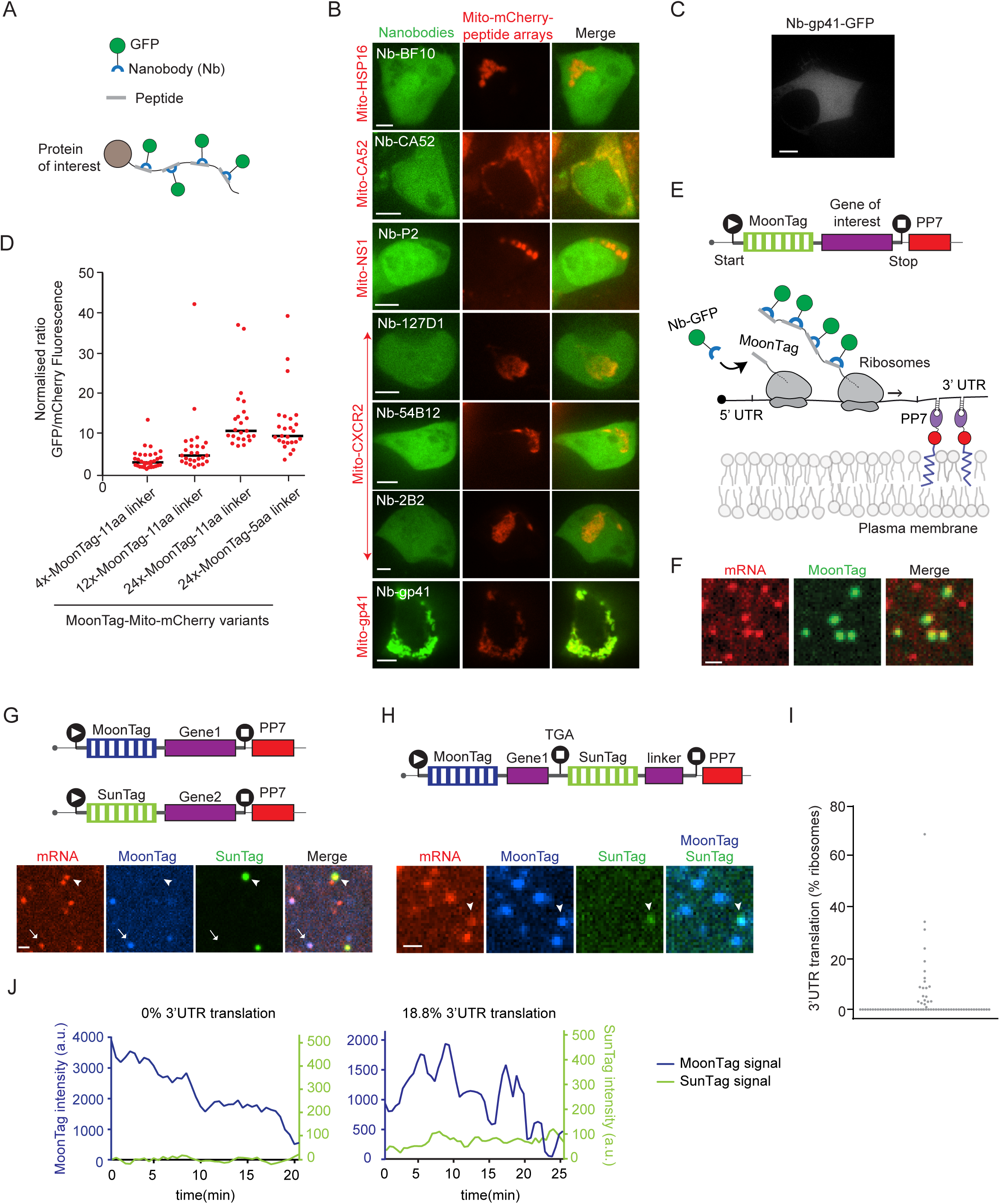
Development of the MoonTag, a new fluorescence labeling system to visualize translation of single mRNAs. A) Schematic representation of the nanobody-peptide labeling system. B) Representative images of HEK293T cells transfected with indicated GFP-labelled nanobodies along with twelve copies of their respective peptide epitopes fused to mCherry and to a protein domain that targets to the outer mitochondrial surface (Mito). Scale bars, 5µm. C) Representative image of a HEK293T cell transfected with MoonTag nanobody-GFP. Scale bar, 10 µm. D) Indicated constructs were transfected in U2OS cells stably expressing GFP-labeled MoonTag nanobody. The GFP:mCherry fluorescence intensity ratio on mitochondria was quantified. Each dot represents a single cell, and lines indicates the average value of all analyzed cells. E) Schematic of mRNA reporter used for translation imaging (top panel). Nascent polypeptide labelling strategy using the MoonTag system (bottom panel). The MoonTag nanobody binds to and fluorescently labels nascent polypeptides, while the PP7 system is used to fluorescently label and tether mRNAs. F) U2OS cells stably expressing MoonTag-Nb-GFP and PCP-mCherry-CAAX expressing the MoonTag translation reporter shown in (E). Representative image of a few translating mRNAs is shown. Scale bar, 1µm. G-H). Schematic of reporters (top) and representative images of Moon/Sun cells expressing indicated reporters (below). G) Arrow heads and arrows indicate SunTag translation signals and MoonTag translation signals, respectively. Scale bar, 1µm. H) Arrow heads indicate read-through translation signal. Scale bar, 1µm. I-J) Moon/Sun cells were transfected with the reporter indicated in (H) and MoonTag and SunTag intensities on single mRNAs were tracked over time. I) Frequency of 3’UTR translation (percentage of ribosomes) was calculated for individual mRNAs. Every dot represents a single mRNA. J) Example dual-color intensity traces of two mRNAs with MoonTag (blue) and SunTag signal (green). Number of experimental repeats and mRNAs analyzed per experiment are listed in Table S1. See also Movie S1.

To determine the binding stoichiometry of the MoonTag nanobody to its peptide array, we created peptide arrays containing 4, 12 or 24 MoonTag peptides, which were fused to Mito-mCherry. The binding stoichiometry of the MoonTag nanobody and peptide array was then determined by quantitatively comparing mCherry and GFP fluorescence near mitochondria (See Methods), which revealed that up to ~10-12 MoonTag nanobodies bind to an array of 24 MoonTag peptides (Fig. 1D), slightly less than was observed for the SunTag (Tanenbaum et al., 2014). When we reduced the linker length from 11 to 5 amino acids, we did not observe a significant change in MoonTag labeling efficiency. Therefore, to reduce the size of the MoonTag peptide array, we used the array with 5 amino acid linkers for further experiments. Fusion of MoonTag peptides to either a histone or a membrane protein resulted in recruitment of the MoonTag nanobody to DNA and the plasma membrane, respectively (Fig. S1A-B), indicating that MoonTag-fused proteins localize correctly to different cellular compartments. To test whether the MoonTag could also be labeled with different fluorophores, we swapped the GFP for a HaloTag (Los et al., 2008), which was labeled with far-red dye, JF646 (Grimm et al., 2015). GFP or HaloTag-based labeling of the MoonTag resulted in a similar recruitment of the MoonTag nanobody to the peptide array (Fig. S1C), providing the possibility to label the SunTag and MoonTag in different colors and therefore the potential to combine both systems in a single cell.

Next, we introduced the MoonTag peptide array in our previously developed translation imaging reporter (Fig. 1E) (Yan et al., 2016). In brief, the MoonTag is inserted upstream of a gene of interest (in this case, the kinesin Kif18b). During translation, the MoonTag peptides are synthesized before the protein of interest and are rapidly bound by the MoonTag nanobody co-translationally. This results in bright fluorescence labeling of the nascent polypeptide, providing a direct readout of translation of single mRNA molecules. Additionally, the reporter mRNA contains 24 binding sites for the PP7 coat protein (PCP) (Chao et al., 2008) in the 3’UTR. Co-expression of PCP-2xmCherry enables fluorescence labeling of the mRNA independently of translation. The PP7 system was also used to tether the mRNAs to the plasma membrane; PCP was fused to a membrane anchor (the C-terminal CAAX sequence), which results in immobilization of mRNAs near the plasma membrane (Fig. 1E). Membrane tethering of mRNAs substantially increases signal-to-noise during imaging and facilitates long-term tracking of individual mRNAs, without detectably altering translation dynamics (Yan et al., 2016).

The MoonTag translation reporter was transfected in human U2OS cells stably expressing MoonTag-Nb-GFP and PCP-2xmCherry-CAAX. Time-lapse imaging was performed at 30s time interval using a spinning disk confocal microscopy in a single z-plane, focused on the bottom plasma membrane. As expected, MoonTag foci could be observed that co-localized with single mRNAs (Fig. 1E-F), indicating active translation of those mRNA molecules. These results demonstrate that the MoonTag system can be applied to label nascent polypeptides and visualize translation of single mRNA molecules in real-time, similar to the SunTag system.

For simultaneous analysis of two different types of mRNAs in single cells, we generated SunTag and MoonTag translation reporters containing different genes and co-expressed these reporter mRNAs in U2OS cells stably expressing SunTag-scFv-GFP, MoonTag-Nb-Halo^JF646^ and PCP-2xmCherry-CAAX (referred to as Moon/Sun cells). Red mRNA foci were observed that co-localized with either SunTag or MoonTag signal, but no mRNAs were observed that contained both signals (Fig. 1G), demonstrating that the SunTag and MoonTag systems are fully orthogonal and can be used together in the same cell to visualize translation of two different mRNAs.

To test whether the SunTag and MoonTag systems could also be combined in a single mRNA, we generated a translation reporter with the MoonTag inserted in the ORF, while the SunTag was inserted downstream of the stop codon to visualize translation of the 3’UTR (Fig. 1H, schematic). Most mRNAs showed strong MoonTag translation signal, but only a small subset of MoonTag-positive mRNAs showed SunTag signal as well (Fig. 1H). The SunTag signal intensity was generally low, and most individual SunTag translation events only lasted a few minutes (Fig. 1H and Movie S1). 3’UTR translation was likely caused by occasional stop codon read-through by individual ribosomes. While translation reinitiation after termination at the MoonTag ORF stop codon is also possible, it is less likely because no AUG start codons were present in the SunTag reading frame downstream of the stop codon. Surprisingly, large variations in the frequencies of 3’UTR translation were observed between different mRNA molecules (Fig. 1I-J). The vast majority of mRNAs (90.4 %) did not show any 3’UTR translation over the time period of imaging (mean track length 16.9 ± 5.2 min (mean ± SD)), while other mRNAs showed continuous 3’UTR translation, indicative of translation by multiple ribosomes (Fig. 1I-J). The differences in the frequency of 3’UTR translation between different mRNAs were not caused by corresponding differences in the translation initiation rate of those mRNAs (Fig. S1D), suggesting that different mRNA molecules may have a distinct susceptibility for stop codon read-through even though these mRNAs were derived from the same gene. Thus, the SunTag and MoonTag systems can be combined in single cells, and even in single mRNAs to visualize complex aspects of mRNA translation with single ribosome sensitivity.

### Development of a translation reading frame reporter that reports on translation start site selection

Alternative translation start site selection is an important form of translational heterogeneity, as the majority of mRNAs contain multiple translation start sites, and translation start site selection can determine both protein sequence and expression levels. We wondered whether the development of the MoonTag would allow real-time observation of translation start site selection. Since the translation start site determines the reading frame of a ribosome, we reasoned that a reporter of translation reading frame could be leveraged to report on translation start site selection. To develop a translation reading frame reporter, we designed a new tag in which MoonTag and SunTag peptides were “mashed” together: they were fused in alternating fashion and positioned in different reading frames. In this design, all SunTag peptides were located in the −1 reading frame with respect to the MoonTag peptides (Fig. 2A). Note that the +1 frame does not contain any SunTag or MoonTag sequences, and is referred to as the ‘blank’ frame. We named this ribosome reading frame reporter the <u>M</u>oonTag and <u>S</u>unTag <u>h</u>ybrid (Mash)Tag (Fig 2A). To adapt the MashTag so it would report on translation start site selection, we designed two versions of the MashTag reporter. Both versions contained 36 copies of the MashTag (devoid of stop codons in all frames), a downstream gene of interest, followed by stop codons in all three frames. As gene of interest, we designed a BFP sequence lacking stop codons in all frames to ensure that the coding sequence length of the MoonTag and the SunTag frame is equal. Finally, 24 PCP binding sites were introduced in the 3’UTR of the MashTag reporter to visualize and tether mRNAs. One version of the MashTag reporter contained an AUG translation start codon in frame with the MoonTag peptides (referred to as the ‘MoonStart’ reporter), while the other contained an AUG in frame with the SunTag peptides (‘SunStart’ reporter) (Fig. 2B-C, schematics). Both AUGs were placed in strong initiation sequence context (a Kozak consensus sequence: GCCACC<u>AUG</u>G) and no other AUG sequences were present in the 5’UTR or MashTag sequence. To visualize translation of the MashTag reporters, we expressed them in the Moon/Sun cells (U2OS cells stably expressing SunTag-scFv-GFP, MoonTag-Nb-Halo^JF646^ and PCP-mCherry-CAAX). During initial attempts to image cells expressing the MashTag reporters, we noticed that at high expression level, the ‘mature’ (i.e. ribosome released) protein encoded in the SunTag frame of the MashTag tended to form protein aggregates. Therefore, the MashTag reporter was expressed from an inducible promoter (a CMV-based Tet-On promoter) to reduce protein synthesis before the onset of imaging, and all cells that showed protein aggregates were excluded from further analyses.

**Figure 2.**
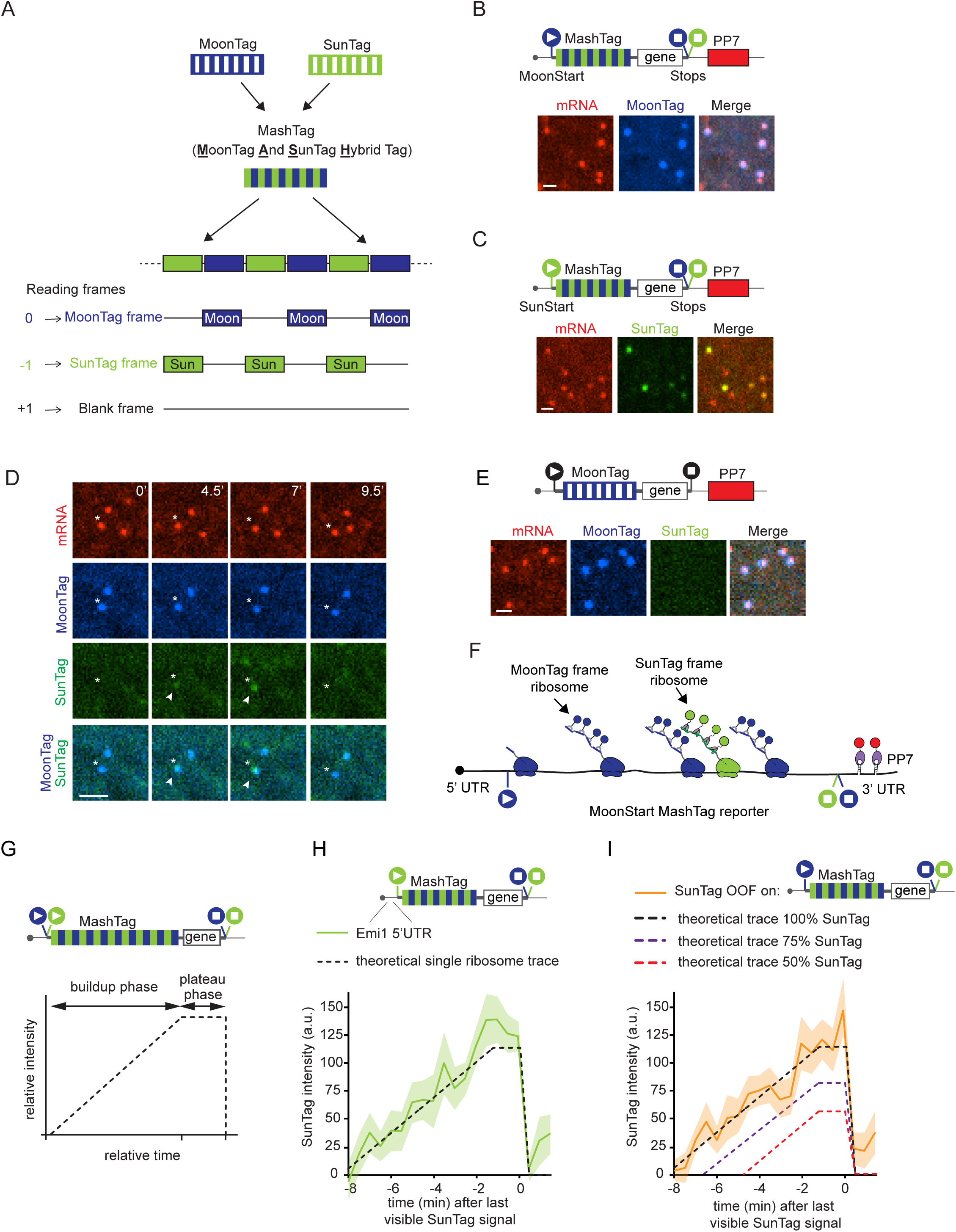
The MashTag: a reading frame sensor to visualize translation start site selection. A) Schematic of the MashTag design. MoonTag (blue) and SunTag peptide (green) repeats are fused in alternating fashion such that translation of the 0 reading frame exclusively results in MoonTag translation signal, whereas translation of the −1 reading frame leads to SunTag translation signal. (B, C, E, G, H, and I) Schematic of MashTag translation reporters (top panels). Filled circles with white triangles represent translation start sites; filled circles with white squares represent translation stop codons. Colors of the filled circles indicate reading frame; blue indicates MoonTag reading frame, while green represent SunTag reading frame. Black start and stop sites in (E) indicate that only a single reading frame is visualized in these reporters. For simplicity, 24xPP7 sites in the 3’UTR are not depicted in the schematics of G-I. B, C, E) (bottom panels): Representative images of translating mRNAs in Moon/Sun cells expressing the indicated translation reporters. Scale bars, 1µm. D) Moon/Sun cells expressing the MashTag translation reporter shown in (B). Different time points of representative mRNAs are shown. Asterisks indicate the mRNA with MoonTag and OOF translation. Arrow heads indicate the OOF SunTag signal. Time is indicated in min. Scale bar, 2µm. F) Schematic of OOF translation on an mRNA of the MashTag reporter shown in (B). G) Schematic illustrating a theoretical intensity trace of a single ribosome translating a MashTag reporter mRNA. The trace contains a buildup phase, during which MashTag is translated and a plateau phase during which the gene of interest is translated. (H-I) Fluorescence intensities of single ribosomes translating the reporter mRNA in the SunTag frame, either when the SunTag was in the main frame (H) or was OOF (I). Intensity traces are aligned at the last time point that contains SunTag signal (i.e. just before translation termination). Solid lines indicate experimentally derived values, shaded areas surrounding solid lines indicate SEM. Black dashed lines (H, I) indicate expected single ribosome intensity trace if all 36 SunTag peptides of the MashTag sequence are translated; red and purple dashed lines (I) indicated expected intensity traces if 50% and 75% of SunTag peptides are translated, respectively. Number of experimental repeats and mRNAs analyzed per experiment are listed in Table S1. See also Movie S2.

MoonStart and SunStart reporters showed predominantly MoonTag and SunTag translation signals, respectively (Fig. 2B, C), indicating that they accurately report on the dominant translation start site. Surprisingly, when analyzing the MoonStart reporter, we observed frequent brief pulses of SunTag signal on mRNA molecules that also showed MoonTag signal (Fig. 2D and Movie S2). These pulses of SunTag signal on the MoonStart reporter could not be explained by background fluorescence, as similar fluorescence signals were not observed on mRNAs containing only the MoonTag (Fig. 2E). The SunTag pulses on the MoonStart reporter mRNAs therefore represent a subset of ribosomes that are translating the MoonStart reporter in the SunTag reading frame, which we will refer to as out-of-frame (OOF) translation (Fig. 2F). Together, these results show that the MashTag reporter can accurately report on the dominant translation start site of an mRNA, and can simultaneously reveal non-canonical, OOF translation events on individual mRNA molecules.

### SunTag frame translation is mainly due to alternative translation start site selection

OOF translation in the MoonStart reporter could either be due to alternative translation start site selection or ribosome frameshifting. Alternative translation start site selection presumably occurs near the 5’end of the MashTag and thus is expected to include most if not all SunTag peptides. In contrast, if OOF translation is caused by ribosome frameshifting on the MashTag reporter, the OOF translation event would contain only a subset of SunTag peptides, reducing both the SunTag fluorescence intensity and duration of the OOF translation event. To differentiate between these scenarios, we wished to compare the fluorescence of OOF translation events to the expected fluorescence signal of a single ribosome translating the entire array of 36 SunTag peptides (referred to as the ‘theoretical single ribosome intensity trace’). The theoretical single ribosome intensity trace contains three distinct phases: 1) a fluorescence intensity buildup phase. During the buildup phase the SunTag peptides are sequentially synthesized and fluorescently labeled by antibodies; 2) a plateau phase when the gene downstream of the MashTag (i.e. the BFP sequence) is translated. During the plateau phase no new SunTag peptides are synthesized, and the fluorescence remains constant; 3) a sudden drop in fluorescence when translation is terminated and the nascent chain is released from the mRNA (Fig. 2G). To determine the duration of the buildup and plateau phases, we calculated the ribosome elongation speed using harringtonine run-off experiments (See Methods), which revealed an elongation speed of 2.9 ± 2.0 codons/s (mean ± SD) (Fig. S2A), similar to our previously determined translation elongation rate in U2OS cells (Yan et al., 2016). Using the nucleotide length of the MashTag and BFP sequences, combined with the experimentally determined translation elongation rate, the duration of the buildup and plateau phases could be calculated (429 sec and 74 sec, respectively). Next, we determined the fluorescence intensity during the plateau phase. The plateau intensity represents the fluorescence intensity of a single, fully synthesized array of 36 SunTag peptides encoded by the MashTag, and was determined to be 110 ± 53 a.u. (mean ± SD) (Fig. S2B, see Methods).

To validate the values for the theoretical single ribosome intensity trace, we directly determined the fluorescence intensity over time of a single ribosome translating the entire 36 repeats of the MashTag reporter in the SunTag frame. To image single translating ribosomes, we introduced the highly repressive 5’UTR of Emi1 into the SunStart reporter, which reduces translation initiation rates by ~50-fold and as a result maximally one ribosome translates an mRNA molecule at the time (Tanenbaum et al., 2015; Yan et al., 2016). Comparison of the theoretical and observed single ribosome intensity traces revealed highly similar traces (Fig. 2H), demonstrating that the theoretical intensity trace accurately represents the fluorescence associated with a single ribosome translating the entire 36 repeats of the MashTag.

We also generated two additional theoretical intensity traces, which represent translation of either 18 or 27 SunTag peptides by a single ribosome, the approximate average number of SunTag peptides that would be translated if SunTag OOF signal was caused by frameshifting on the MoonStart reporter (Fig. 2I) (See Methods). We then analyzed the SunTag fluorescence intensity traces of OOF translation events on the MoonStart reporter and compared them to either the trace containing all 36 SunTag peptides, or the traces containing 18 or 27 peptides. This comparison revealed that the intensity profile of single OOF translation events was very similar to the theoretical intensity trace of 36 SunTag peptides (Fig. 2I), suggesting that OOF translation is predominantly caused by alternative start site selection near the 5’ end of the mRNA.

To confirm that SunTag translation was mainly caused by alternative translation start site selection and not by frameshifting, fluorescence intensities of mature SunTag proteins were measured that were synthesized either from the MoonStart reporter (OOF translation) or from the SunStart reporter (which contains all 36 SunTag repeats in frame with the AUG start codon). Fluorescence intensities of mature proteins originating from MoonStart and SunStart reporters were very similar (Fig. S2C), confirming that frameshifting was not a major cause of OOF translation signal. Note that OOF fluorescence could, in theory, also be explained by frameshifting that occurs exclusively at a unique sequence near the 5’ end of the MashTag. However, this is unlikely, as the 5’ region of MashTag sequence does not contain any sequences known to induce frameshifting. Additionally, the nucleotide sequence of the MashTag is quite repetitive, so any frameshifting sequence in one of the first MashTag repeats is likely present multiple times in the MashTag, and thus not unique to the 5’ end. Together, these analyses indicate that most of the OOF SunTag translation events are caused by alternative start site selection. Therefore, our MashTag reporter can be used to study translation start site selection kinetics and variability.

### A computational pipeline to quantitatively interpret fluorescence signals on translating mRNAs

To understand the heterogeneity and dynamics of translation initiation at both canonical and alternative start sites, it is key to extract quantitative information about the frequency and timing of both types of initiation events. To determine the precise number of ribosomes, as well as the time of translation initiation of each ribosome in the both the MoonTag and SunTag frame, fluorescence intensities of MoonTag and SunTag need to be measured over time for each mRNA molecule. To facilitate intensity measurements, we developed an automated analysis package in Matlab with a graphical user interface (GUI) (“TransTrack”, which we will freely distribute upon publication). TransTrack enables simultaneous mRNA tracking and fluorescence intensity measurements in multiple colors, and generates fluorescence intensity traces for both SunTag and MoonTag frames for each mRNA as output.

The next step in determining the timing of individual translation initiation events, is the conversion of SunTag and MoonTag fluorescence signals to the number of ribosomes at each time point. The SunTag or MoonTag fluorescence at each time point of an intensity trace represents one or more ribosomes translating the mRNA (Fig. 3A). To determine the precise number of ribosomes that contribute to the fluorescence of a translating mRNA, we made use of the theoretical fluorescence intensity profile of a single ribosome. To achieve this for both the SunTag and MoonTag frames, we generated a theoretical intensity trace of a single ribosome for the MoonTag frame, similar to the theoretical SunTag trace (also see Fig. 2G, See Methods). By positioning one or more theoretical single ribosome intensity traces along the time-line of an experimentally observed translation site intensity trace, and summing up their intensity profiles at each time point, the experimentally observed intensity trace of an mRNA translated by multiple ribosomes can be reconstructed *in silico* (Fig. 3A). We developed an iterative stochastic modeling approach to identify the number and position of translation initiation events which generated the best fit with the experimental data (Fig. 3B, See Methods). Reproducible high quality fits were obtained after 1000 iterations of optimization (Fig. 3B and S3D-F) (See Methods). Using this fitting approach, we could determine the number of ribosomes in both frames at each time point (bottom panel of Fig. 3B) and the moment of translation initiation of each ribosome in either frame (indicated by triangles underneath the x-axes in Fig. 3B). To validate our tracking and fitting algorithms, we generated three control reporters: one reporter containing only SunTag peptides, one containing only MoonTag peptides, and one reporter containing both SunTag and MoonTag peptides that were placed in the same reading frame (‘Moon-SunTag’ reporter). As expected, when SunTag-or MoonTag-only reporters were analyzed, ribosomes were detected almost exclusively in the SunTag and MoonTag frames, respectively (Fig. 3C). Furthermore, the Moon-SunTag reporter showed a narrow distribution in the ratio of SunTag and MoonTag signals, centering close to 50% (Fig. 3C), confirming the accuracy of our analysis pipeline.

**Figure 3.**
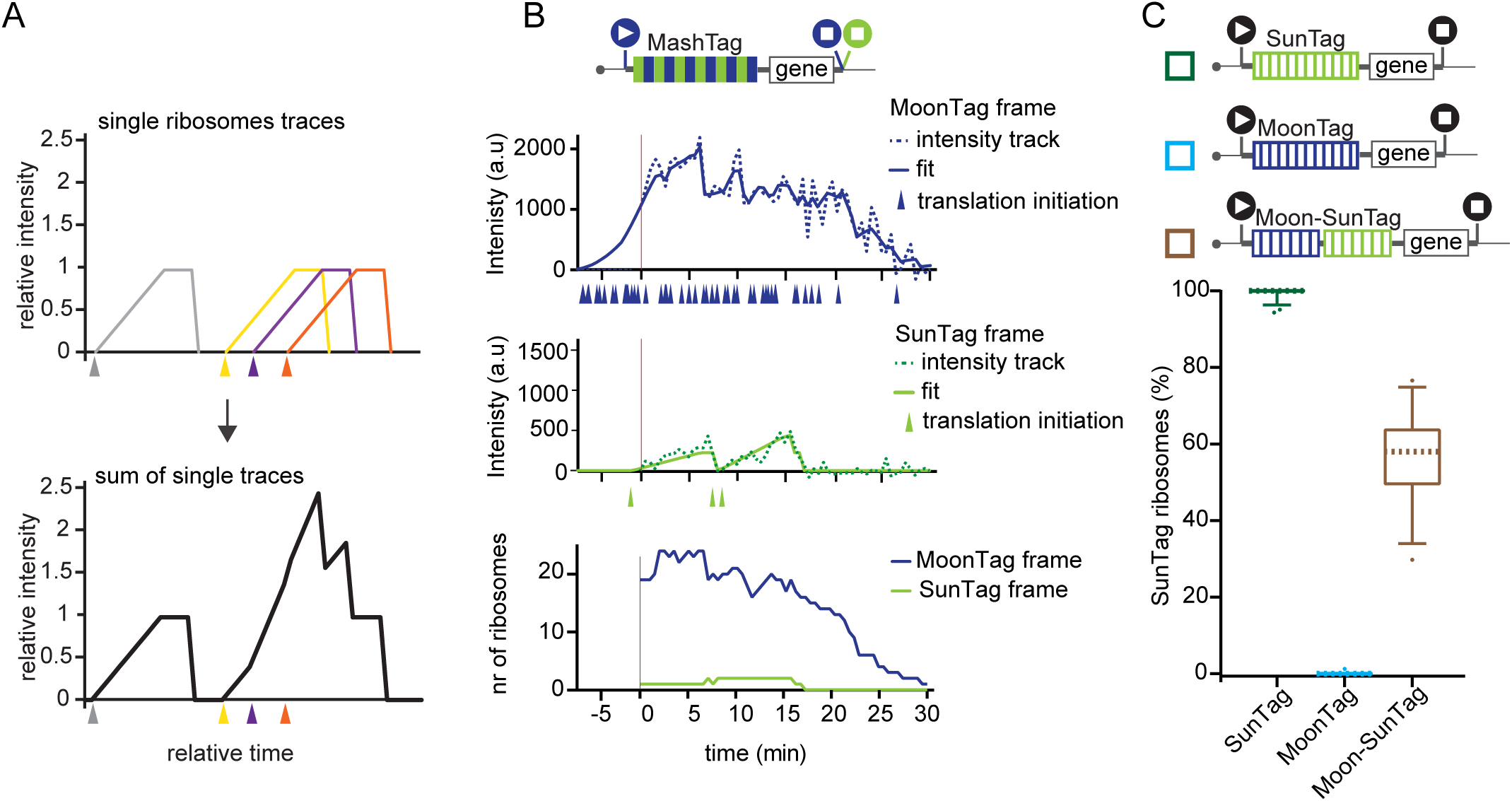
A computational pipeline to quantitatively interpret fluorescence signals. A) Schematic of determining number of ribosomes and moment of initiation per ribosomes based on an intensity trace. Each single ribosome leads to a theoretical single ribosome trace (top panel), which is observed as a cumulative trace of all single theoretical traces (bottom panel). Each color represents a single ribosome. Triangles indicate translation initiation moment of corresponding single ribosomes. B-C) Schematic reporters: filled circles with white triangles represent translation start sites; filled circles with white squares represent translation stop codons. Colors of the filled circles indicate reading frame; blue indicates MoonTag reading frame, while green represents SunTag reading frame. Black start and stop sites in (C) indicate that only a single reading frame contains MoonTag or SunTag peptides in these reporters. For simplicity, 24xPP7 sites in the 3’UTR are not depicted in the schematics. B) An example dual-color intensity trace of a single MashTag mRNA with MoonTag (blue; top panel) and SunTag (green; middle panel) signal. Dashed lines indicate experimentally observed intensities, solid lines display the optimal fit. Colored triangles below the x-axes of top and middle graphs represent translation initiation events. Bottom panel shows ribosome occupancy per reading frame over time as determined by fitting of the same mRNAs as top to panels. Red vertical line indicates the first time point of the trace. C) Boxplots indicate the relative percentage of ribosomes translating the SunTag frame (relative to the sum of MoonTag and SunTag frame) on single mRNAs of the indicated reporter mRNAs. The number of ribosomes translating either the MoonTag or SunTag reading frame was determined based on fitting of ribosomes to intensity tracks (as described in B). Dashed line represents median value, box indicates 25-75% range, and whiskers indicated 5-95% range. Number of experimental repeats and mRNAs analyzed per experiment are listed in Table S1.

### Analysis of translation start site selection dynamics and heterogeneity

To determine the frequency of OOF translation on the MoonStart reporter, intensity traces were generated for 85 mRNA molecules that contained detectable translation in either reading frame, and the number of ribosomes translating either reading frame was determined for each mRNA. Traces had a duration of 26 ± 6 min (mean ± SD) and contained 38 ± 30 (mean ± SD) translation initiation events. Most mRNAs were strongly translated in the MoonTag frame; 87 % mRNAs had an initiation rate of > 0.5 ribosomes per min in the MoonTag frame. The majority of mRNA molecules (66%, 56 out of 85) showed both SunTag and MoonTag translation events, even within the relatively short time-window of our observations, indicating that multiple translation start sites are used intermittently on most mRNA molecules. Surprisingly, we observed widespread variability in the frequency of OOF translation, ranging from 0% to 100% of the ribosomes (median, 7%) (Fig. 4A-B). Two possible explanations could account for the observed differences in OOF translation frequency on different mRNA molecules. First, it is possible that translation start site selection is stochastic, and some mRNAs have more OOF translation than others by chance. In this model, the probability of initiating translation at each potential start site is identical for every mRNA molecule. Alternatively, the probability of alternative start site selection may be distinct for different mRNA molecules. To distinguish between these possibilities, we performed statistical analyses, which revealed that for 25% (21 of 85) of mRNAs start site usage frequency deviated significantly from the population (Fig. 4C) (p < 0.01, binomial test, See Methods). These results indicate that the probability of using alternative start sites is distinct among different mRNAs and show that different mRNA molecules originating from a single gene can be heterogeneous with respect to translation start site usage.

**Figure 4.**
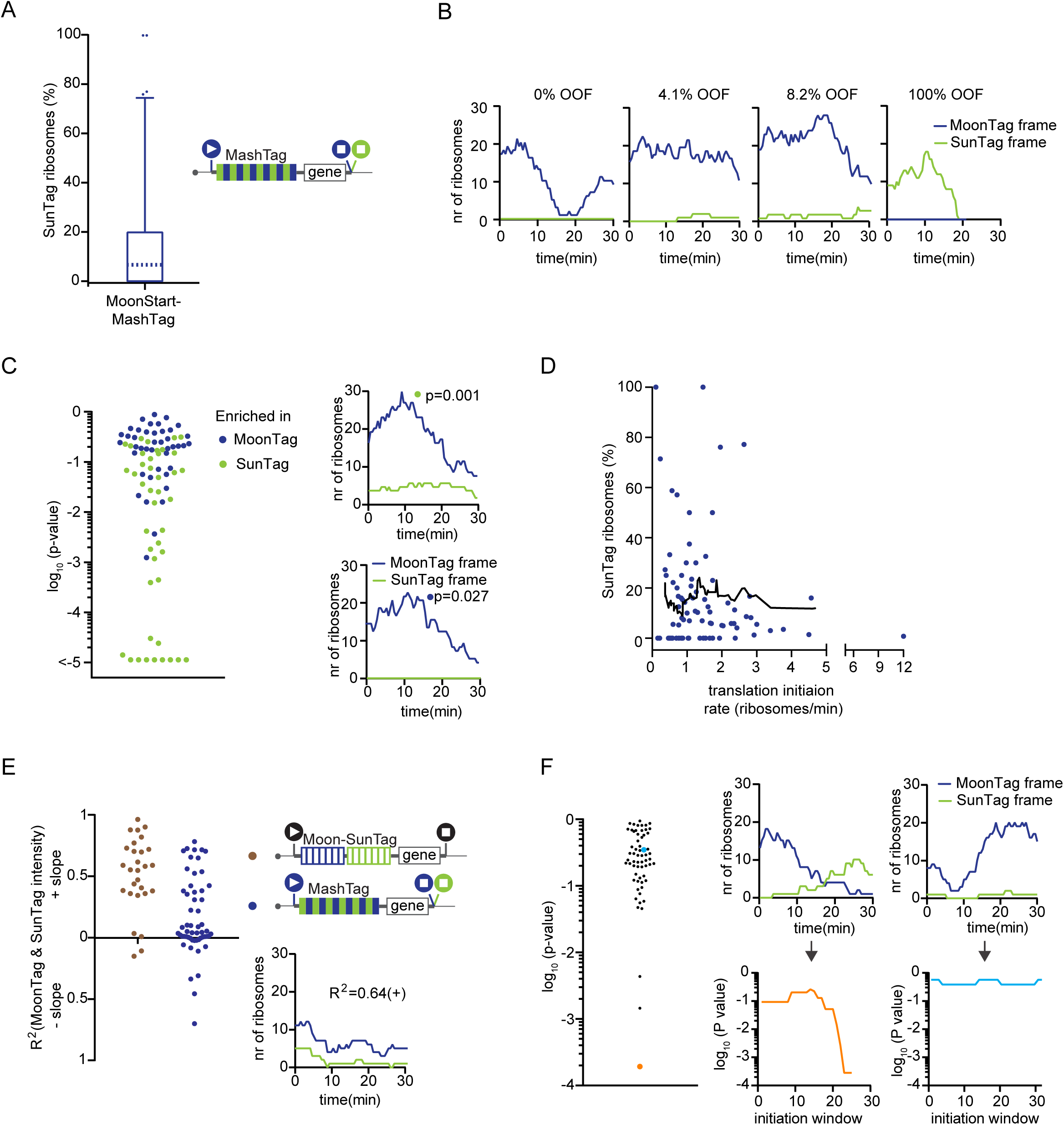
Heterogeneity in translation start site selection among different mRNA molecules. A, E) Schematics: filled circles with white triangles represent translation start sites; filled circles with white squares represent translation stop codons. Colors of the filled circles indicate reading frame; blue indicates MoonTag, while green represents SunTag reading frame. Black start and stop sites in (E) indicate that only a single reading frame is visualized in these reporters. For simplicity, 24xPP7 sites in the 3’UTR are not depicted in the schematics. A-F) Indicated reporters were transfected into Moon/Sun cells and MoonTag and SunTag intensities on single mRNAs were tracked over time. (A) Boxplot indicates the relative percentage of ribosomes translating the SunTag frame (relative to the sum of the MoonTag and SunTag frame) on single mRNAs. Dashed line represents median value, box indicates 25-75% range, and whiskers indicated 5-95% range. B) Example graphs of ribosomes in each reading frame over time for four mRNAs of the reporter indicated in (A). Percentages represent fraction of SunTag ribosomes on the mRNAs. C) Distribution of p-values after binomial testing for enrichment of SunTag or MoonTag ribosomes on mRNAs compared to the median fraction of SunTag ribosomes as determined in (A). Every dot represents a single mRNA (top graph). Example graphs of ribosome numbers over time in each reading frame for individual mRNAs (bottom graphs). P-values for example traces are indicated. D) Correlation between overall translation initiation rate and relative SunTag frame translation frequency for individual mRNAs of the reporter indicated in (A). Every dot represents a single mRNA and the line depicts moving average over 15 mRNAs (See Methods). E) Linear regression analysis of MoonTag and SunTag intensities for indicated reporter mRNAs (top graph). Example trace of one mRNA is shown (bottom graphs) with indicated R^2^ value. F) Sliding window analysis (see Fig. S4 for details) of initiation events in MoonTag and SunTag reading frames on mRNAs of the reporter indicated in (A). Every dot depicts the strongest p-value of a single mRNA (left graph). Example traces of ribosomes in each reading frame over time (top right graphs) and corresponding sliding window p-values (bottom right graphs). Dots and traces related to the same mRNA are labeled in the same color. Number of experimental repeats and mRNAs analyzed per experiment are listed in Table S1.

To test whether alternative start site selection frequency depends on the overall translation initiation rate of an mRNA (i.e. the sum of MoonTag and SunTag initiation rates), we compared for each mRNA molecule the frequency of OOF translation with the overall translation efficiency. The OOF frequency was similar over a range of translation initiation rates (Fig. 4D), demonstrating that OOF translation does not depend on the overall translation efficiency. Next, we asked whether translation initiation rates in the MoonTag and SunTag frames were correlated over time. To determine whether initiation at different sites is co-regulated, we performed linear regression analysis on the intensities of SunTag and MoonTag translation signals for all time points of an mRNA. As a positive control, the level of correlation between SunTag and MoonTag fluorescence over time was determined in the Moon-SunTag reporter, which showed a strong positive correlation, as expected (Fig. 4E). Analysis of SunTag and MoonTag fluorescence on the MoonStart reporter also revealed a positive correlation between translation in both reading frames for many mRNAs, albeit not as strong as the Moon-SunTag reporter; 38% of MoonStart mRNAs (21/56) showed a positive correlation (R^2^ > 0.2) (Fig. 4E). The positive correlation between MoonTag and SunTag translation initiation events over time may be explained by temporal fluctuations (i.e. bursting) in the rate of ribosome recruitment to the mRNA, which could affect the initiation rate at all start sites.

A smaller subset (7%) of mRNAs showed a negative correlation between SunTag and MoonTag signal (R^2^ < −0.2) (Fig. 4E), suggesting that distinct start sites could also be regulated differentially over time. To investigate independent regulation of different start sites in more detail, the relative frequency of SunTag and MoonTag initiation was determined over shorter periods of time to detect “bursts” in the usage of particular translation start sites. The SunTag and MoonTag translation initiation frequencies were determined in a sliding window of 10 sequential translation initiation events, and the relative initiation frequencies for each window were compared to the average translation frequencies of both frames of the entire trace (Fig S4A-B; see Methods). We then calculated the probability of observing the relative SunTag and MoonTag initiation frequency of each window and determined the lowest window p-value of each mRNA (Fig. 4F). This sliding window analysis revealed that the majority of temporal fluctuations in the relative frequency of SunTag and MoonTag translation can be explained by chance, indicating that on individual mRNAs start site selection is largely stochastic. However, on a small number of mRNAs (5%; 3/63) a statistically significant (p < 0.01) change in translation start site selection was observed during the time of observation (Fig. 4F). While the observed frequency of such bursts in translation start site usage in our dataset was relatively low, our average observation time of individual mRNAs was only 26 min, representing ~10% of the life-time of a typical mRNA (Schwanhäusser et al., 2009), so the fraction of mRNAs that undergoes changes in translation start site usage may be far higher.

### Alternative translation start site selection can occur on near-cognate start sites both upstream and downstream of the AUG start codon

Since the MoonStart reporter does not contain any AUG start sites in the SunTag frame, SunTag translation must initiate on near-cognate start codons, which could be located upstream or downstream of the MoonTag AUG start site (Fig. 5A). Downstream start sites could be encountered by ribosomes after ‘leaky’ scanning (i.e. scanning over the start site without initiation) of the MoonTag start site. To test whether leaky scanning of the MoonTag AUG start codon results in OOF translation on the MoonStart reporter mRNAs, a second AUG start codon was inserted into the mRNA downstream of the MoonTag AUG. Introduction of additional start sites in the MoonTag or blank frame significantly reduced the number of translation initiation events in the SunTag frame (p < 0.01 and p < 0.001, respectively) (Fig. 5B). Addition of a start site in the SunTag frame downstream of the MoonTag start site slightly increased the SunTag translation signal, although this effect was not significant (p=0.14). However, introduction of additional start sites in the blank frame between the MoonTag and the newly introduced SunTag start site did significantly decrease initiation in the SunTag frame (p < 0.05). Together, these results show that leaky scanning of the MoonTag start site followed by downstream initiation on a near-cognate start codon in the SunTag frame contributes to OOF translation on the MoonStart reporter.

**Figure 5.**
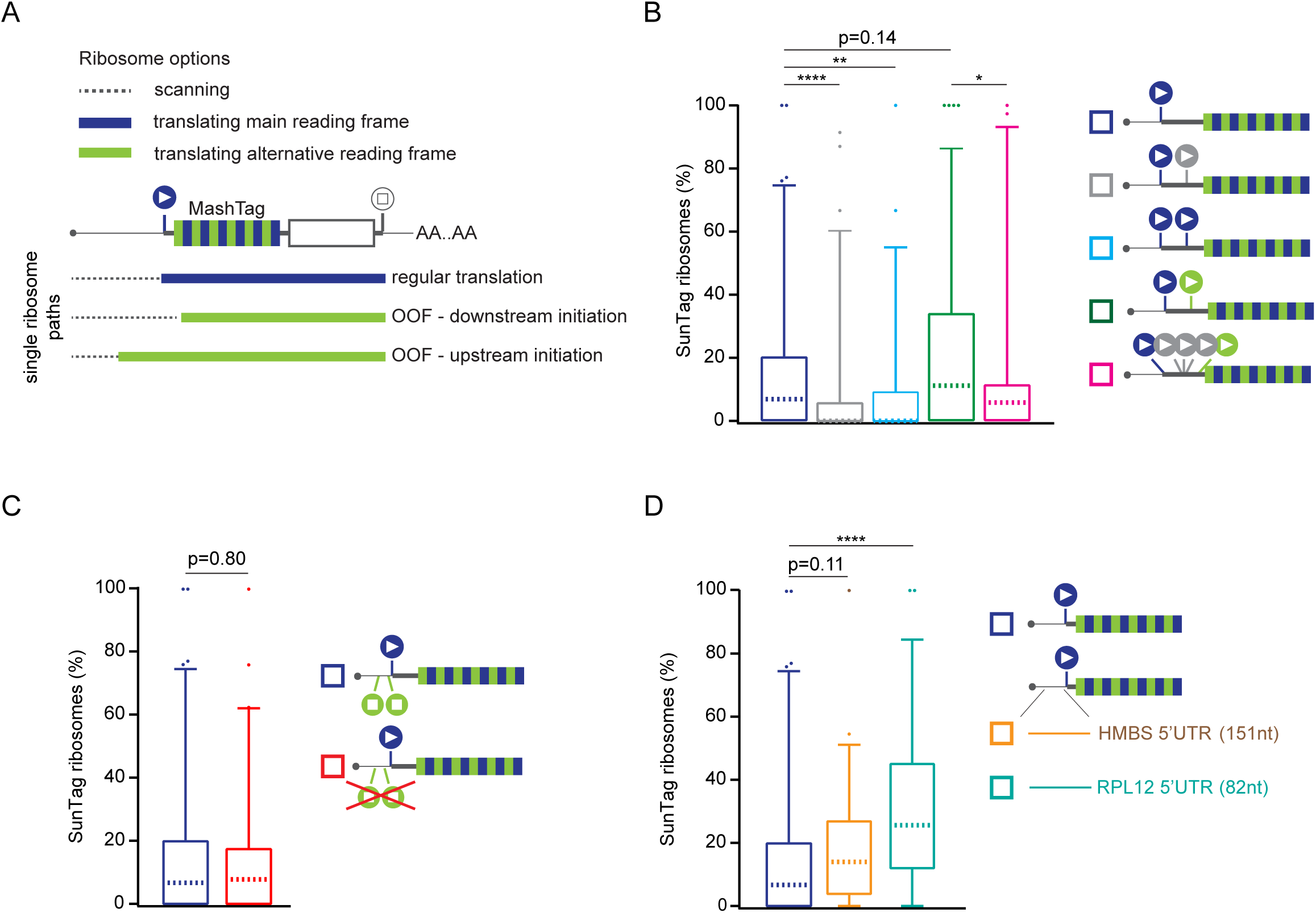
Alternative start site selection contributes to OOF translation. A) Schematic of different possible translation paths of individual ribosomes on a MashTag mRNA. (B-D) Schematic reporter: filled circles with white triangles represent translation start sites; filled circles with white squares represent translation stop codons. Colors of the filled circles indicate reading frame; blue indicates MoonTag, green indicates SunTag reading frame. For simplicity, reporter schematics only indicate 5’ region of the mRNA. B-D) Indicated reporters were transfected into Moon/Sun cells and MoonTag and SunTag intensities were tracked over time on single mRNAs. Boxplots indicate the relative percentage of ribosomes translating the SunTag frame (relative to the sum of the MoonTag and SunTag frame) on single mRNAs. P-values are based on two-tailed Mann-Whitney test; * p<0.05; ** p<0.01; *** p<0.001; **** p<0.0001. For comparison, data indicated with dark blue (B-D) is re-plotted from Fig. 4A. Dashed line represents median value, box indicates 25-75% range, and whiskers indicated 5-95% range. Number of experimental repeats and mRNAs analyzed per experiment are listed in Table S1.

Next, we wished to examine the role of upstream start site selection in OOF translation of the MoonStart reporter. The MoonStart reporter contained two stop codons in the SunTag frame upstream of the MoonTag AUG, which would prevent upstream translation initiation from generation SunTag signal. Therefore, these two stop codons were removed from the reporter (MoonStart ΔSunStops) (Fig. 5C, schematic). Removal of the stop codons did not significantly increase the level of SunTag translation (Fig. 5C), suggesting that upstream initiation in the SunTag reading frame does not strongly contribute to OOF translation on this reporter mRNA.

Rocaglamide A (RocA), an inhibitor of the translation initiation factor eIF4A, was recently shown to stimulate upstream translation initiation (Iwasaki et al., 2016). Since we did not observe any upstream translation initiation on the MoonStart reporter, we tested whether upstream initiation could be induced by RocA treatment. For these experiments, the MoonStart ΔSunStops reporter was used to ensure that upstream initiation would result in translation of the SunTag. 0.5 μM RocA treatment resulted in a 37% reduction in overall translation, consistent with inhibition of a key translation initiation factor (Fig. S5A). However, the relative fraction of ribosomes initiating translation in the SunTag frame markedly increased from 8.7% to 21.4 % (median values, p < 0.01), while treatment with DMSO as a control did not significantly alter the ratio in SunTag and MoonTag translation (8.7% vs 8.0%, median values, p = 0.33) (Fig. S5B). These analyses show that upstream translation start site selection can also result in OOF translation, and confirm the previous finding that RocA can stimulate upstream translation initiation.

To access upstream translation initiation on endogenous 5’UTR sequences, two additional MashTag reporters were generated that contained the 5’UTRs of two genes, HMBS and RPL12 (151 and 82 nucleotides in length, respectively), both of which lack start and stop codons in the SunTag frame. While the HMBS 5’UTR reporter showed a similar OOF translation frequency as the MoonStart reporter, introduction of the RPL12 5’UTR into the reporter resulted in a substantially increased OOF translation frequency (median, 25.6% vs 7.0%; p < 0.01) (Fig. 5D). Interestingly, the overall initiation rate of the RPL12 5’UTR reporter was also reduced by 28% compared to the MoonStart reporter (Fig. S5C) (p < 0.001), indicating that the RPL12 5’UTR contains strong translation regulatory elements that results in OOF translation These results demonstrate that extensive upstream translation initiation occurs on natural 5’UTR sequences, suggesting that alternative start site selection might be a widespread phenomenon on endogenous mRNAs.

### A real-time sensor to visualize translation of uORF-containing mRNAs

uORFs are present in thousands of mRNAs and generally represses translation of the downstream (main) ORF (Calvo et al., 2009; Chew et al., 2016; Johnstone et al., 2016; Wethmar, 2014). Ribosomes that translate a uORF can dissociate from the mRNA after translation termination at the uORF stop codon, thus preventing translation of the downstream ORF. Translation of the main ORF can occur either through uORF skipping (i.e. leaky scanning of the uORF start site), or through translation reinitiation at the downstream ORF after translation termination at the uORF stop codon. While a previous study has used the SunTag system to visualize translation of a protein coding ORF downstream of a uORF (Wang et al., 2016a), real-time visualization of multiple translation paths (e.g. uORF translation vs uORF skipping) of a uORF-containing mRNA was not feasible, and therefore the frequency and heterogeneity in path selection by different ribosomes could not be assessed. To determine uORF translation, uORF skipping, and translation reinitiation in real-time on single mRNAs, we generated a single molecule uORF sensor using the MashTag (Fig. 6A). The uORF sensor is based on the MoonStart reporter and thus contains an AUG start codon in the MoonTag frame. Upstream of the MoonTag AUG, the reporter contains a short uORF (48 nucleotides; similar to the median human uORF length (Calvo et al., 2009)). The uORF start codon was placed in the blank frame, so initiation at the uORF start site could not result in MoonTag or SunTag fluorescence. A third AUG codon was inserted into the coding sequence of the uORF, and was placed in frame with the SunTag (Fig. 6A). In this reporter, SunTag signal reports on leaky scanning of the uORF start codon, while a MoonTag signal reflects translation reinitiation after uORF translation. Ribosomes that dissociate from the mRNA after uORF translation are not *directly* observed, but can be inferred from the decrease in MashTag translation (i.e. MoonTag + SunTag) upon introduction of the uORF into the reporter.

**Figure 6.**
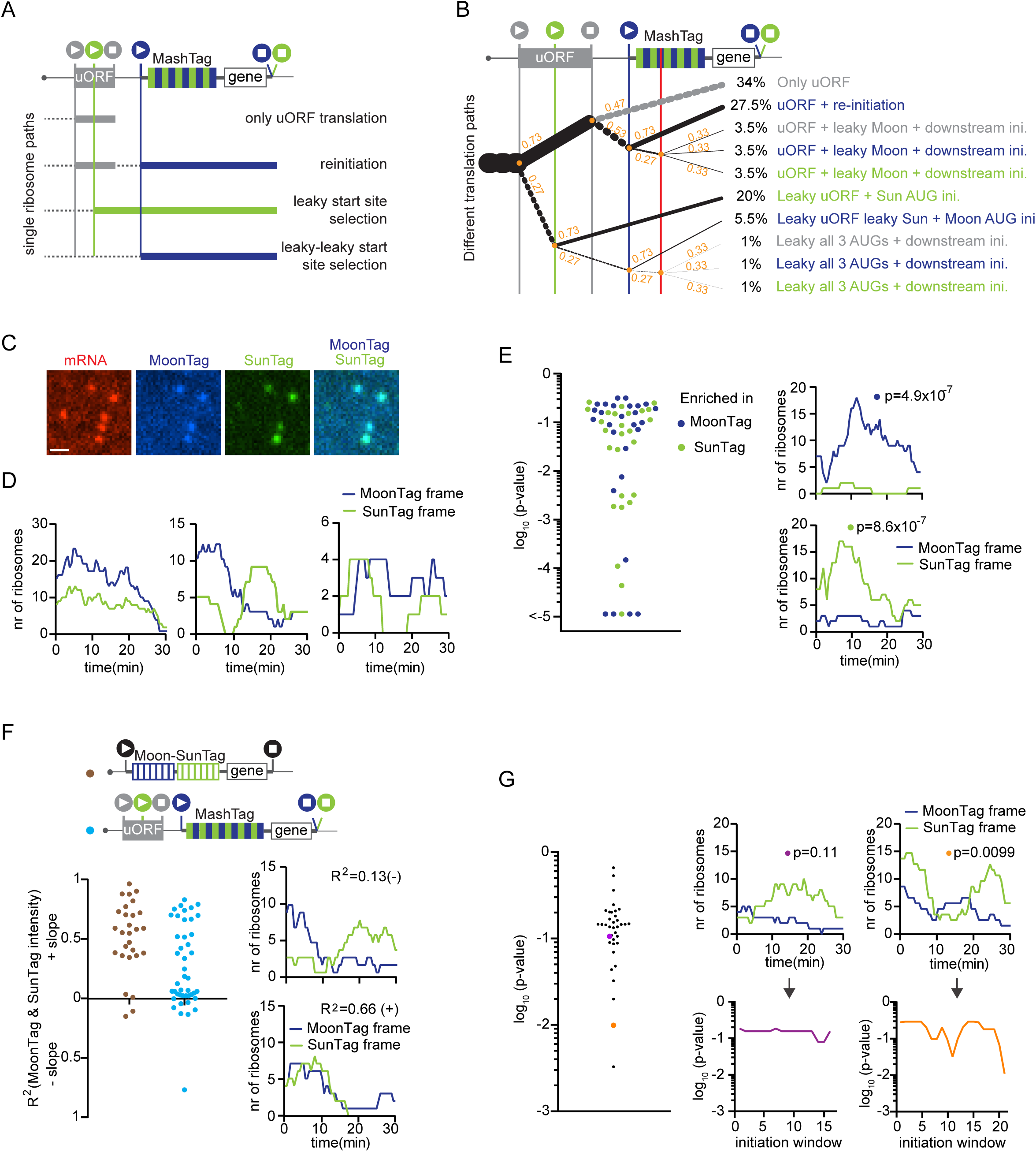
A single molecule uORF sensor based on the MashTag. A, B and F) Schematic reporters: filled circles with white triangles represent translation start sites; filled circles with white squares represent translation stop codons. Colors of the filled circles indicate reading frame; blue indicates MoonTag, green indicates SunTag, and grey indicates blank reading frame. Black start and stop sites in (F) indicate that only a single reading frame is visualized in these reporters. For simplicity, 24xPP7 sites in the 3’UTR are not depicted in the schematics. A) Schematic of different possible translation paths of individual ribosomes on a uORF-MashTag mRNA. B) Fraction of ribosomes undergoing each translation path. Thickness of line reflects the fraction of ribosomes undergoing the path. Dashed lines indicate ribosome scanning; solid dashed lines indicate translation. Dashed grey line indicates termination after uORF translation. Orange numbers at branch points indicate fraction of remaining ribosomes that follow each path. Red line indicates non-canonical start sites in all three frames in the MashTag. Colors of path descriptions correspond to frame in which the MashTag is translated. C-G) MashTag reporters were transfected into Moon/Sun cells and MoonTag and SunTag intensities on single mRNAs were tracked over time. C, D) Representative images (C) and example graphs (D) of translating mRNAs in a cell expressing the reporter indicated in (A). Scale bar, 1µm. E) Distribution of p-values (binomial test) for enrichment of SunTag or MoonTag ribosomes on mRNAs of reporter indicated in (A) were analyzed. Ratio of ribosomes translating the SunTag and MoonTag for each individual mRNA was compared to the median fraction of SunTag ribosomes, as determined in Fig. S6D. Every dot represents a single mRNA (left graph). Example graphs of two mRNAs are shown with indicated p-values (right graphs). F) Linear regression analysis of MoonTag and SunTag intensities for indicated reporter mRNAs (left graph). For comparison, data indicated in brown is replotted from Fig. 4E. Example graphs of two mRNAs are shown with indicated R^2^ values (right graphs). G) Sliding window analysis (see Fig. S4 for analysis details) of initiation events in MoonTag and SunTag reading frames on mRNAs of reporter indicated in (A). Every dot depicts the strongest p-value of a single mRNA (left graph). Example traces of ribosomes in each reading frame over time (top right graphs) and corresponding sliding window p-values (bottom right graphs). Dots and tracks related to the same mRNA are labeled with the same color. Number of experimental repeats and mRNAs analyzed per experiment are listed in Table S1.

Based on the translation rates in both SunTag and MoonTag frames, and based on the overall reduction of translation of the MashTag upon insertion of the uORF, we could estimate the frequency of usage of all translation paths along the uORF reporter (see Methods); 34% of ribosomes translate the uORF and do not reinitiate, 27.5% of ribosomes translate the uORF and reinitiate on the downstream MoonTag start site, 20% of ribosomes skip the uORF AUG through leaky scanning and initiate on the SunTag AUG, and the remaining 19% of ribosome follow more complex paths (Fig. 6B). We also swapped SunTag and MoonTag start sites, such that SunTag signal reports on translation reinitiation and MoonTag signal reports on uORF AUG leaky scanning, which resulted in similar values for uORF translation and reinitiation (Fig. S6A). To experimentally confirm our calculations on the usage of different translation paths, we removed the uORF stop codon, extending the uORF coding sequence beyond the MoonTag AUG start site (Fig. S6B-D). In this reporter, MoonTag signal is no longer caused by translation reinitiation. Based on our calculations, we predicted that this would result in 83% reduction in MoonTag signal. Indeed, we found a strong (79%) reduction in MoonTag signal after removal of the uORF stop codon, whereas the SunTag translation rate was unaffected (p=0.75) (Fig. S6B-C) (see Methods). This result quantitatively confirms our calculations of the different translation paths, and also confirms that the large majority of MoonTag translation is due to translation reinitiation (Fig. 6B). Together, these results reveal that the MashTag-based uORF sensor can provide a quantitative readout of all possible paths that ribosomes can take along a uORF-containing mRNA molecule.

Next, translation of individual uORF-containing mRNA molecules was examined in more detail. Most mRNAs (44/53) contained both SunTag and MoonTag signal (Fig. 6C, D), demonstrating that ribosomes following different paths along the mRNA (e.g. uORF skipping and translation reinitiation) co-exist on most mRNA molecules. However, the relative frequency of the different translation paths varied between different mRNA molecules. A subset of mRNAs (15/53) showed a significantly greater fraction of translation in either the MoonTag or the SunTag frame than expected based on the total population of mRNAs (Fig. 6E, p>0.01), demonstrating that the probability of uORF skipping and translation reinitiation is variable among different mRNA molecules.

When examining the precise moment of translation initiation of ribosomes translating either the SunTag or MoonTag reading frames, a strong temporal correlation between SunTag and MoonTag translation signals was observed on many mRNAs (Fig. 6F). As discussed before, this correlation is likely caused by the burst-like behavior of translation initiation, which results in fluctuations in the number of ribosomes that are loaded on the mRNA over time. While the majority of mRNAs showed a positive correlation over time between SunTag and MoonTag translation, a more detailed analysis using the sliding window approach (see Fig. S4A-B and Methods) revealed that a subset of mRNA molecules (6/37, p< 0.05) showed statistically significant bursts of either translation reinitiation or uORF skipping (Fig. 6G), suggesting that uORF translation may be dynamically regulated over time on individual mRNAs. Bursts in translation start site selection did not take place simultaneously on all mRNAs in the same cell, suggesting that the regulation of uORF translation does not occur in a cell-wide manner, but rather at the level of individual mRNA molecules. Together, these results provide the first real-time observations of uORF translation, uORF skipping and translation reinitiation, offer a quantitative assessment of all the paths that ribosomes take along the 5’UTR of a uORF-containing mRNA and provide a powerful new assay to study mechanisms of translation regulation by uORFs.

## Discussion

In this study, we develop a new system to fluorescently label nascent polypeptides, the MoonTag, which can be used to visualize mRNA translation and is orthogonal to our previously developed SunTag system. Combining the SunTag and MoonTag systems in single cells and single mRNAs provides new opportunities to investigate complex dynamics of mRNA translation. Here, we combine the MoonTag and SunTag systems to generate the MashTag, a real-time single molecule reporter of translation reading frame usage, which we leverage to visualize translation start site selection by individual ribosomes. Using the MashTag system, we find a striking heterogeneity in translation start site selection among different genes, different mRNA molecules originating from the same gene, and different time points in the life of a single mRNA. Thus, the MoonTag and MashTag provide a powerful new set of tools for studying translation dynamics and heterogeneity.

### Applications of dual-color single molecule translation imaging

The SunTag system is limited to analysis of a single type of mRNA, and a single read-out per mRNA. However, many aspects of translation are more complex, requiring knowledge of multiple parameters of translation for a complete understanding of the process. Therefore, we developed the MoonTag, an orthogonal translation imaging system. Expression of SunTag and MoonTag mRNAs in the same cell enables the direct comparison between two different types of mRNAs, for examples between different genes or different mRNA isoforms. It may be feasible to visualize even more than two mRNA species when combining the SunTag and MoonTag; by generating additional peptide arrays that consist of MoonTag and SunTag peptides (in the same frame) mixed together at different ratios, it could become possible to distinguish numerous different mRNA species in the same cell. Adding a third orthogonal nascent chain labeling system, for example the ‘spaghetti monster’ (Viswanathan et al., 2015), would further increase the possible number of mRNA species that can simultaneously be analyzed.

The SunTag and MoonTag systems can also be combined in single mRNAs to interrogate complex aspects of translation. In this study, we show that dual color translation imaging can be used to assess translation of the 3’UTR, translation start site selection, and the dynamics of uORFs translation. In the accompanying paper (Lyon et al.), a similar dual-color translation imaging approach is developed, and the authors investigate the kinetics of ribosome frameshifting on a viral RNA sequence. While the two studies investigate different biological processes, both studies uncover a high degree of heterogeneity among different mRNA molecules, and striking temporal fluctuations in translation dynamics of individual mRNAs (e.g. bursts in translation initiation and frameshifting). As similar conclusions are drawn for such diverse molecular processes, it is likely that widespread translational heterogeneity is the norm, rather than an exception, highlighting the importance of studying translational heterogeneity and dynamics in detail. The dual-color translation imaging approach will be an important tool to unravel the prevalence, kinetics, and molecular mechanisms of such translational heterogeneity. In many cases, one of the two colors (e.g. the MoonTag) is used to measure canonical translation, while the other system reports on the alternative pathway (e.g. frameshifted ribosomes). This approach allows for the analysis of the precise number of ribosomes that follow each path on each mRNA, which is not feasible with a single color translation imaging system. The ability to quantitatively determine different translation pathways simultaneously for a single mRNA was, for example, essential to distinguish bursts in ribosome recruitment to the mRNA from start site specific initiation bursts.

To facilitate quantitative analysis of dual-color translation imaging data, we have also developed two new software packages, the first (TransTrack) for tracking and three color intensity measurements of single mRNAs, and a second package to convert fluorescence intensities into ribosome numbers and translation initiation times. These packages, which will be made freely available through Github (URL upon publication), and will be invaluable for future approaches using multi-color translation imaging, as well as for many single color translation imaging approaches.

### Mechanisms of translation start site selection heterogeneity

Using the MoonStart reporter, we found that, overall, ~7% of ribosomes show OOF (i.e. SunTag frame) translation. This value likely represents a lower limit for endogenous genes, as; 1) our MashTag system only reports on translation of one alternative reading frame and OOF translation probably also takes place in the other alternative (blank) frame; 2) the MoonStart reporter contains a very strong translation start site, limiting leaky scanning and downstream initiation. In contrast, genome-wide analysis of all translation start sites revealed that many genes have suboptimal start site context (Noderer et al., 2014); 3) The MashTag reporter contains a very short, unstructured 5’UTR lacking regulatory elements or additional AUG sequences, limiting upstream start site selection. Endogenous 5’UTRs can be far more complex and therefore could result in a substantially higher upstream initiation rate. Indeed, introducing the endogenous 5’UTR sequence of RPL12 significantly increased OOF translation. Together, these findings suggest that alternative start site selection and OOF translation is likely a widespread phenomenon on endogenous mRNAs.

A subset of mRNA molecules in our analysis (~25%) showed a significantly higher likelihood of translation initiation on alternative start sites as compared to the bulk of mRNAs. There are several possible explanations for the altered probability of translation initiation on different mRNAs. First, differences in transcription start site usage create mRNAs with different 5’UTRs, which, in turn, could affect translation start site usage, and TSS usage is known to be highly variable in mammalian cells (Forrest et al., 2014). Second, RNA modifications, specific RNA structures or binding of regulatory proteins may alter the probability that translation is initiated on a given start site. Indeed, the mRNA modification m6A was recently implicated in translation start site selection (Zhou et al., 2018), and mRNA structures can also bias translation initiation site selection in yeast (Guenther et al., 2018). While differences in nucleotide sequence would result in a permanent difference in translation start site usage, RNA modifications, RNA structures and RBP-dependent regulation could be dynamically regulated to alter start site usage over time. For a number of mRNAs, we indeed observed a significant change in relative start site usage over time, demonstrating that start site selection can be dynamically regulated for single mRNAs. Identifying regulatory mechanisms that shape start site usage will be an important future goal, and the MashTag system will be a valuable tool for investigating such mechanisms.

For many mRNAs, the average usage of different translation start sites was similar. Nonetheless, the timing of translation initiation and the precise order of initiation events in different reading frames was unique for each mRNA molecule, which likely reflects the inherent stochasticity in start site selection by individual ribosomes. Our results also revealed that the frequency of translation initiation at MoonTag and SunTag start sites was positively correlated over time. We have previously shown that the translation rate is not constant over time on individual mRNAs, but rather shows a burst-like behavior (Yan et al., 2016). The fact that multiple translation start sites show correlated bursting, suggests that the burst-like behavior of translation originates upstream of translation start site selection, likely at the step of 43S ribosome recruitment to the mRNA. In summary, while translation start site selection by individual ribosomes appears mostly stochastic, the probability of usage of individual start sites is under tight control, probably both transcriptionally and post-transcriptionally.

Translation of uORF-containing mRNAs showed many similar characteristics as translation of mRNAs with a single AUG start codon: 1) most mRNAs contained multiple intermittently-used translation start sites; 2) the selection of a translation start site by individual ribosomes appeared stochastic; 3) usage of different start sites tended to correlate over time; 4) a substantial fraction of mRNA molecules (28%) showed a distinct translation start site usage pattern compared to the bulk of mRNAs; 5) temporal burst in uORF translation, uORF skipping and/or translation reinitiation could be observed. These results suggest that the dynamics and heterogeneity of start site selection are inherent properties of translation and are likely valid for many types of mRNAs.

### Consequences of widespread alternative translation start site selection

Pervasive variability in start site selection likely has major implications for cellular function. In-frame alternative start site selection results in N-terminally extended or truncated proteins. Many genes encode N-terminal localization signals, like mitochondrial or ER targeting signals, required for proper protein localization. Production of proteins lacking such N-terminal signals would affect protein localization and functionality. In contrast, OOF translation initiation results in proteins with a completely different amino acid sequence, which are likely misfolded and non-functional. Synthesis of such OOF protein products not only would waste cellular energy, but could cause a considerable proteotoxic stress to the cell as well. The costs of OOF translation may be partially negated, because ribosomes that initiate translation at an OOF start site are likely to encounter an early termination codon, which can trigger nonsense-mediated mRNA decay (Lykke-Andersen and Jensen, 2015). mRNAs that have a high probability of OOF translation may therefore be rapidly degraded, reducing the amount of OOF translation products that are synthesized. While widespread alternative translation initiation can be problematic, the high degree of flexibility in translation start site selection can also be exploited by the cell. For example, it enables different types of post-transcriptional gene regulation (e.g. uORF-dependent translational control), and it may be important for regulated changes in N-terminal protein sequences as well. An important future question is whether the extensive OOF translation that we observed on different mRNAs is functionally important for the cell, or rather reflect errors in translation start site selection.

The frequent occurrence of initiation downstream of the main ORF AUG start codon observed in our experiments also has important implications for genetic engineering using CRISPR/Cas9. To obtain a gene ‘knockout’, a frameshift mutation is often generated downstream of the AUG of the protein coding ORF. Our results reveal that a substantial fraction of ribosomes fail to initiate at the protein coding AUG start site. If these ribosomes initiate translation downstream of the frameshift mutation on an in-frame start codon, significant expression of an N-terminally truncated protein can be expected. Therefore, leaky scanning of the main start site needs to be taken into consideration when designing a knockout strategy.

In summary, by developing the MoonTag system we have expanded the toolbox to study translation dynamics of single mRNAs. The combination of the MoonTag and SunTag systems offers two independent readouts of translation and will provide an invaluable tool to elucidate complex translation dynamics.

## Acknowledgements

We would like to thank members of the Tanenbaum lab for helpful discussions. We would also like to thank Merlijn Staps for initial work on the computational pipeline and Tim Hoek for critically reading the manuscript. This work was financially supported by an ERC starting grant (ERC-STG 677936-RNAREG), grants from the Netherlands Organization for Scientific Research (NWO) (ALWOP.290 and NWO/016.VIDI.189.005) and this work was supported by Howard Hughes Medical Institute through an International Research Scholar grant to MET (HHMI/IRS 55008747) and at Janelia Research Campus (JBG and LDL). MET was also financially supported by the Oncode Institute.

## Author contributions

SB, DK, and MET conceived of the project. JBG and LDL provided reagents. BMPV and SS developed and optimized the software. SB and DK performed all the experiments. SB, DK, and BMPV analyzed the data. SB, DK, and MET prepared the figures and MET wrote the manuscript with input from SB and DK.

## Material and methods

### 1. General methods

#### 1.1 Plasmids

The following nanobody sequences were obtained and ordered as G-blocks from IDT:

- Nb-BF10 (Srivastava et al., 2013);
- Nb-CA52 (Smolarek et al., 2010);
- Nb-2B2, Nb-127D1, and Nb-54B12 (Bradley et al., 2015);
- Nb-P2 (Fatima et al., 2014);
- Nb-gp41 (Lutje Hulsik et al., 2013).

All peptide array sequences were ordered from Genewiz. To design the MashTag, the following considerations were taken into account: 1) each repeat of the SunTag or MoonTag in the MashTag had to encode the same SunTag or MoonTag amino acid sequence; 2) no AUG start codons or stop codons (TGA, TAA, or TAG) were introduced in any reading frame; 3) different codons were used for the same amino acid sequence in different copies of the SunTag and MoonTag peptides to introduce nucleotide sequence variation between individual repeats; 4) all sites for restriction enzymes were removed. After generation of a MashTag containing plasmid, the size of the MashTag was checked by enzyme digestion, and the 5’ and 3’ ends were sequence verified. Because of difficulties in sequencing due to the repetitive nature of the MashTag, the middle part of the MashTag was not sequence verified for all plasmids.

#### 1.2 Lentiviral infections and cell line generation

To produce lentiviruses, HEK293T cells were infected with the lentivirus plasmid of interest and lentiviral packaging vectors ps.Pax and p.MD2 using PEI (Polyenthylenimine). One day after transfection, the medium was replaced with fresh medium. Virus-containing medium was collected 3 days after transfection. To infect U2OS cells with lentivirus, cells were seeded 24h before infection and grown to ~60% confluency at moment of infection. The supernatant of the HEK293T cells containing the lentivirus was added to the U2OS cells. U2OS cells were spin-infected for 90-120 minutes at 2000 rpm at 25°C. After spin-infection, the medium was replaced with fresh medium and cells were cultured for at least 2 days before any further analysis or processing. Where applicable, cells were FACS-sorted as single cells in 96-well plates to generate monoclonal cell lines.

#### 1.3 Cell culture

Human U2OS cells and HEK293T (ATCC) were grown in DMEM (4.5g/L glucose, Gibco) containing 5% fetal bovine serum (Sigma-Aldrich) and 1% penicillin/streptomycin (Gibco). Cells were grown at 37°C and with 5% CO2.

#### 1.4 Microscopy

##### 1.4.1 Single molecule translation imaging

For translation imaging experiments, all imaging was done using U2OS cell lines stably expressing TetR (for inducible expression), PP7-2xmCherry-CAAX, and either MoonTag-Nb-GFP or MoonTag-Nb-Halo^JF646^ and scFv-sfGFP. Cells were seeded in glass bottom 96-wells plates (Matriplates, Brooks) at 15-20% confluency 2 days before imaging. DNA plasmids encoding reporter mRNAs were transfected 1 day prior to imaging using Fugene (Promega) and for MashTag imaging experiments, a BFP-encoding plasmid was co-transfected (DNA ratio 1:1), which was used for initial identification of transfected cells. One hour prior to imaging, medium was replaced with CO_2_-independent pre-warmed L15/Leibovitz’s (Thermo Fisher) containing 50nM Halo^JF6464^. After Halo incubation for 1h at 37°C, the cells were briefly rinsed twice with L15/Leibovitz’s medium and washed once with L15/Leibovitz’s for 15 minutes. Doxycycline (1 μg/ml) was added 15-20 minutes before start of imaging to induce transcription of the reporter. To select cells for imaging, approximately 50 positions were first selected based on BFP signal (the co-transfection marker). From this selection, approximately 10 positions were chosen for time-lapse imaging based on the presence of translation sites and the absence of protein aggregates. For time-lapse imaging, images were acquired at 30s interval with 500ms exposure times for 30 minutes. A single Z-plane was imaged, which focused on the bottom plasma membrane of the cells. Images were acquired using a Nikon TI inverted microscope with perfect focus system equipped with a Yokagawa CSU-X1 spinning disc, a 100x 1.49 NA objective and an iXon Ultra 897 EM-CCD camera (Andor) using Micro-Manager Software (Edelstein et al., 2010) or NIS Elements Software.

##### 1.4.2 Screening of antibody-peptide pairs

For screening of Nanobody-peptide pairs, seven different nanobodies fused to GFP, were cotransfected with their respective mito-mCherry-peptide arrays in HEK293T cells and analyzed for co-localization at mitochondria using a spinning disc confocal microscope.

### 2. mRNA tracking and intensity measurements

#### 2.1 Tracking of single mRNAs to obtain MoonTag and SunTag intensities

Maximally 10 mRNAs were tracked per cell. mRNAs were selected randomly, regardless of the type of translation signal (either MoonTag or SunTag). To measure the intensity of fluorescence on mRNAs over time, we generated a semi-automated translation spot tracking in MatLab called TransTrack. Upon publication, TransTrack will be made available, including documentation, through Github.

#### 2.2 Correction of fluorescence intensities

Bleach correction was performed in TransTrack. In brief, to correct for photobleaching during the imaging, the fluorescence intensity of the entire field of view was determined at each time point of the movie. The fluorescence intensity over time was fit with an exponential decay distribution to determine the bleaching rate, and this rate was used to correct all fluorescence images.

We also found that cells with higher expression of the MoonTag-nanobody showed on average higher intensities of MoonTag translation sites (Fig. S3A). Therefore, MoonTag translation site intensities were normalized to total cell MoonTag intensities. As SunTag translation site intensities poorly correlated with total cell intensities, no further correction was performed for the SunTag signal (Fig. S3B).

### 3. A computational pipeline to quantitatively interpret fluorescence signals

#### 3.1 Determine theoretical single ribosome trace

##### 3.1.1 Determining translation elongation rates on MashTag mRNAs

The elongation rates of ribosomes on individual mRNAs was determined by harringtonine run-off experiments as described previously (Yan et al., 2016). In brief, the translation inhibitor harringtonine (3ug/ml) is added to cells expressing the SunStart reporter, which prevents new ribosomes from translating the reporter, and allows ribosomes that are already in the translation elongation phase to continue translating until they reach the stop codon. As ribosomes terminate one-by-one, the SunTag signal of the terminating ribosome dissociates from the mRNA, resulting in a gradual reduction of GFP fluorescence on the mRNA until all ribosomes have terminated translation. The fluorescence decrease was tracked for each mRNA and normalized to the average intensity of the 3 time points before drug administration. Translation elongation rates were then calculated based on the slope of the GFP intensity trace, as described previously (Yan et al., 2016).

##### 3.1.2 Determining the plateau intensity of ribosomes translating the SunTag frame

The plateau intensity of a single ribosome translating the MashTag reporter (Fig 2G), is equal to the intensity of a mature MashTag protein. To determine the intensity of mature proteins translated in the SunTag frame, mature proteins were tethered to the plasma membrane by encoding a C-terminal prenylation sequence (CAAX) in the reporter in the SunTag frame. This reporter was transfected into Moon/Sun cells used for MashTag translation imaging. No doxycycline was added in these experiments to keep the amount of mature protein to a minimum and thereby enable single molecule imaging of mature proteins. Using identical imaging parameters for GFP, as those used during translation imaging, images were acquired of cells containing mature membrane-tethered SunTag protein. For 15 foci (i.e. mature proteins) per cell the intensity was determined. For local background correction, the intensity within a ROI of the same size was determined in a region next to the foci. The mean intensity of 24xMashTag foci in the SunTag frame was 73.5 ± 35.7 a.u. (mean ± SD). For each reporter, the plateau intensity was calculated based on the number of SunTag repeats, therefore this value was correct to 110.3 a.u., as the MashTag used for translation imaging contained 36 instead of 24 repeats (see Table S2 for other reporters).

##### 3.1.3 Determining the plateau intensity of ribosomes translating the MoonTag frame

The plateau intensity of a ribosome translating the MashTag in the MoonTag frame could not be determined with the approach described above, as single mature proteins encoded in the MoonTag frame could not be reliably detected. As an alternative approach, the MoonTag translation site intensity of MoonStart mRNAs was determined (Fig. S3C). For this, only mRNAs were included that contained exclusively MoonTag signal. The mean MoonTag intensity on MoonStart mRNAs was 1354.13 ± 739.65 a.u. (mean ± SD). This fluorescence signal originated from multiple ribosomes. Therefore, to determine the approximate MoonTag intensity associated with a single ribosome, the average number of ribosomes translating a MashTag mRNA was determined. Since the MashTag reporter contained the same promoter, 5’UTR and start codon as our previously described SunTag reporter (Yan et al., 2016), and both reporters had similar translation elongation rates, we assumed that both reporters had similar ribosome densities. Based on the previously determined inter-ribosomal distance on the SunTag reporter (Yan et al., 2016), the average number of ribosomes on a MashTag mRNA could be calculated, which was 17.8. Based on the MoonTag translation site intensity and the number of ribosomes per mRNA, the average MoonTag intensity associated with a single ribosome on the MashTag reporter could be calculated, which was 76.07 a.u. Since only the ribosomes located downstream of the MashTag (i.e. on the BFP sequence) were associated with the full 36 copies of the MoonTag, a correction needed to be applied to obtain the intensity of a 36xMoonTag protein (i.e. the plateau intensity). After applying this correction (as described previously (Yan et al., 2016)), the plateau intensity of a ribosome translating the 36xMashTag in the MoonTag frame was calculated to be 132.7 ± 72 (mean ± SD), and the intensity of a single MoonTag frame encoded MashTag repeat was derived (3.69 a.u.). The combination of this single repeat intensity value and the number of repeats in a reporter was used to determine the plateau intensity of a single ribosome on each reporter (Table S2).

#### 3.2 Validate theoretical single ribosome trace

##### 3.2.1 Comparison between theoretical and experimentally observed intensity traces of single ribosomes

Based on the combination of the translation elongation rate, the plateau intensities and the length of the MashTag and downstream BFP, the theoretical single ribosome traces in either the MoonTag or the SunTag frame could be calculated (Table S2).

To determine the theoretical single ribosome intensity trace for a ribosome translating a MoonStart mRNA in the SunTag frame, two different scenarios were considered: 1) ribosomes could translate the MashTag reporter in the SunTag frame from the start of the coding sequence until the stop codon, or 2) ribosomes could start translating the coding sequence in the MoonTag frame and frameshift into the SunTag frame partway through translation elongation. In the first scenario, the SunTag translation intensity trace on the MoonStart reporter would be similar to the SunTag intensity trace on the SunStart reporter (i.e. containing all 36 SunTag peptides). In the second scenario, fewer SunTag peptides would be translated, resulting in a shorter duration of the intensity trace and a lower plateau intensity. Assuming frameshifting occurs at a random position on the MashTag sequence, frameshifting would occur on average at the end of 18^th^ MashTag repeat (halfway through translating the 36xMashTag). Therefore, the expected intensity trace according to the second model was calculated based on translation of the remaining 18 SunTag repeats. Frameshifting events that occur in the last few repeats of the MashTag may result in a very weak signal, which could evade detection and bias frameshifted translation events towards early frameshifting events that contain more SunTag peptides. Therefore, we also calculated the theoretical intensity trace assuming on average 27 SunTag peptides were translated, which corresponds to a detection limit of 18 SunTag peptides, well within our detection range (Yan et al., 2016).

To generate intensity traces of single ribosomes translating the SunTag frame, both for the Emi1-5’UTR-SunStart reporter and the MoonStart reporter, translation events were manually selected that likely represented single ribosome translation events. For this, all translation events were selected that contained SunTag signal in at least 3 consecutive time points (1.5 min) and at most 16 consecutive time-points (8 min). The 8 min threshold was chosen based on the expected duration of a single ribosome translating the MashTag reporters. All translation events that matched these criteria were aligned at the moment of GFP disappearance (i.e. translation termination) and the GFP intensity of the preceding 8 min was determined. All traces that contained a second translation event within the 8 min preceding time-period, or traces starting in the first 8 min of the movie were removed from the analysis. Average intensities and SEMs of all translation events included in the analysis were then calculated.

##### 3.2.2 Intensity measurements of mature proteins

Experimental settings and analysis of the MoonStart-36xMashTag-CAAX and SunStart-36xMashTag-CAAX (Note that the CAAX motif was in the SunTag frame in both cases) were similar to the experiments described in section 3.1.2 to determine the plateau intensity of the SunTag frame translating ribosome, except that higher laser powers and shorter exposure times were used to facilitate detection of single mature proteins.

#### 3.3 Fitting of raw intensity traces to determine the number and timing of translation initiation events per reading frame

A computational model was developed to determine the number of ribosomes translating an mRNA and the exact time at which single ribosomes initiated translation. The model reconstituted a raw intensity trace by positioning one or more single ribosome translation events along the trace, using the theoretical intensity profile of a single translating ribosome. The sum intensity for each time point of all theoretical single ribosome intensity traces is calculated and compared to the raw intensity trace. By optimizing the number and the position of ribosomes in an iterative fashion the model achieved the optimal fit to the data.

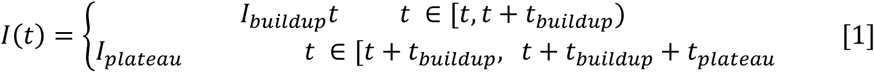

Formula [1] describes the intensity profile of a single translating ribosome, which initiates translation at time t (also see Fig. 2G). I_buildup_, t_buildup_ and t_plateau_ are constant values per reporter, and are dependent on the length and number of MashTag repeats of each reporter, as described in sections 3.1 (Table S2). The raw intensity traces were reassembled *in silico* by fitting the sum of one or more theoretical intensity traces of individual ribosomes that initiate translation at time points t=t_i_ [2]. The *in silico* fitting of ribosomes was performed on the SunTag and MoonTag signals independently and the output from both signals per mRNA were combined after the fitting.

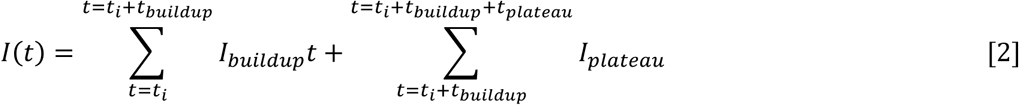

To obtain an initial estimate of the number of ribosomes translating an mRNA based on the raw intensity trace, the area under the curve (AUC) of the entire intensity trace was divided by the AUC of the intensity trace of a single translating ribosome. If the AUC of an mRNA did not exceed the expected AUC of at least one translating ribosome, no further fitting of ribosomes to the intensity trace was performed and the number of ribosomes in the raw intensity trace was determined to be 0. If the raw intensity curve was determined to contain one or more translation events, the model estimated the number of translation events in each raw intensity trace based on the AUC values. These translation events were then distributed along the raw intensity trace with a probability that is weighted by the intensity of the trace at each time point. The sum of all the positioned single ribosome intensity traces was then determined, and the sum intensity trace was compared with the raw intensity trace to determine a goodness of fit, which was defined as the root mean square error (RMSE) between the fit and the data.

After initial placement of translation events, the number of ribosomes and their time of initiation (i.e. their relative position along the trace) were altered according to the parameters shown in Table 1, resulting in a new fit. If the RMSE between the new fit and the data was lower (i.e. the fit improved), the new positions of translation events were used as a starting point for the next iteration. If the RMSE did not improve, the previous positions were used again as a starting point for the next iteration. Of note, if the best fit was achieved with a trace containing no ribosomes, the number of ribosomes was considered 0, even if the AUC of the total intensity trace exceeded the AUC of a single ribosome. The process of re-positioning translation events an accepting or rejecting the new positions was repeated for 1000 iterations to obtain a good fit. A limit of 1000 iterations was selected, because minimal improvements in the fit were achieved with additional iterations (Fig. S3D-F). Since the fitting process is stochastic, multiple runs of the algorithm could result in different outcomes. The algorithm was therefore run 10 independent times for each intensity trace to check for variations in position of translation initiation events and the final RMSE, and the run with the best fit was used to generate the final fit. Note that the 10 runs generally resulted in very similar fits (Fig. S3D), demonstrating the robustness of this approach.

**Table 1:** Parameters used to fit intensity trace to fitting trace.

| Parameter | Value |
| --- | --- |
| Probability to locate a ribosome at time point $t$ in first iteration | $I(t) / \sum_0^{t_{end}} I(t)$ |
| Number of iterations | 1000 |
| Probability to add or remove a ribosome | 0.1 |
| Probability to relocate the time point at which the ribosome initiates | 0.3 |
| Relocation distance (sec) (i.e. repositioning a ribosome along the trace) | The distance to move a translation event is randomly drawn from a normal distribution with $\mu=0$ sec. and $\sigma=25$ sec. |

Many mRNAs already contained SunTag and/or MoonTag signal at the first time point of the intensity trace. However, translation initiation of the ribosomes present on the mRNA at the first time point of the movie took place before the start of the trace, preventing proper fitting. To overcome this limitation, hypothetical time points were added before the start of the intensity trace. The number of added time points depended on the t_buildup_ per reporter. Addition of extra time points allowed the model to position initiation events on the trace prior to the start of image acquisition and, hence, to generate a signal that resembles the intensity at t=0. In the initial placement of ribosomes during the first iteration, the probability of positioning an initiating ribosome before t=0 was equal to the maximal probability to position an initiation event at any time point of the data. The hypothetical time points were not included in calculating the RMSE, as there was no raw intensity measurement that could be compared to the fit during this period.

As output, the model generated an overview containing the following information for both the MoonTag and the SunTag signal on each mRNA: 1) the curves of the best fit and their RMSEs, 2) the total number of ribosomes per reading frame on the mRNA, 3) the initiation timing of each ribosome. The initiation time and the duration of translation of a single ribosome could then be used to calculate for each time point the number of ribosomes per reading frame on the mRNA.

### 4. Quantitative image analysis

#### 4.1 Stoichiometry of MoonTag nanobody binding to MoonTag peptide arrays

The number of MoonTag nanobodies that could bind to a peptide array was determined as described previously (Tanenbaum et al., 2014).

#### 4.2 Calculating translation initiation rates

To determine translation initiation rates of individual mRNAs, the total number of ribosomes translating both SunTag and MoonTag frames was calculated for each mRNA, as described in section 3.3. Initiation rates in each frame were then calculated by dividing the total number of initiating ribosomes by the duration of the trace. Overall initiation rates were based on the sum of all ribosomes translating the MoonTag and the SunTag frames. Ribosomes translating the blank frame were not included in the calculation of initiation rates, as they could not be determined.

#### 4.3 Quantifying 3’UTR translation

In the 3’UTR translation reporter, translation of the coding sequence results in MoonTag signal, while translation of the 3’UTR results in SunTag signal. To determine the number of ribosomes translating the 3’UTR for each mRNA, the number of ribosomes translating both MoonTag and SunTag was determined as described in section 3.3. Because the 3’UTR translation reporter had different numbers of SunTag and MoonTag repeats and the coding sequence was different in length than the MashTag reporter, the theoretical single ribosome traces for translation of both the MoonTag and SunTag were calculated independently (Table S2). The theoretical single ribosome intensity trace of the SunTag was based on a buildup time of 191 s and a plateau time of 72 s, which correspond to the time to translate 24x SunTag repeats and the downstream linker sequence (BFP), respectively. The single ribosome trace of the MoonTag was based on a buildup time of 159.5s and a plateau time of 286s, which correspond to the time to translate 24xMoonTag repeats and the downstream sequence (kif18b). However, if a ribosome reads through the stop codon (which is likely the predominant mechanisms of 3’UTR translation in this reporter), the plateau phase for MoonTag translation is longer (572 s instead of 286 s), as the sequence that is translated downstream of the MoonTag signal now includes the SunTag and BFP as well. A readthrough ribosome therefore gives rise to more total fluorescence. To correct for this, a correction factor was calculated (0.78) based on the fold difference in the AUC of a single ribosome translating the MoonTag and kif18b sequence compared to a ribosome translating the MoonTag, kif18b, SunTag and BFP. To calculate the frequency of 3’UTR translation, the number of SunTag ribosomes was divided by the number of MoonTag ribosomes, after correction.

#### 4.4 Calculating fraction of ribosomes per translation path on the uORF reporter

Translation of a uORF-containing mRNA can result in various different translation paths. The options of single ribosomes were envisioned as a roadmap with a chance to initiate translation or terminate on each start site or stop site encountered by a ribosome, leading to 10 different paths (also see Fig. 6B):

1. Initiation on uORF AUG; termination on uORF; dissociation from mRNA.
2. Initiation on uORF AUG; termination on uORF stop codon; reinitiation of scanning; initiation on MoonTag frame AUG; translation of MashTag in MoonTag frame.
3. Initiation on uORF AUG; termination on uORF stop codon; reinitiation of scanning; skipping of MoonTag frame AUG; downstream initiation in blank frame; translation of MashTag in blank frame.
4. Initiation on uORF AUG; termination on uORF stop codon; reinitiation of scanning; skipping of MoonTag frame AUG; downstream initiation in MoonTag frame; translation of MashTag in MoonTag frame.
5. Initiation on uORF AUG; termination on uORF stop codon; reinitiation of scanning; skipping of MoonTag frame AUG; downstream initiation in SunTag frame; translation of MashTag in SunTag frame.
6. Skipping of uORF AUG; initiation on SunTag frame AUG; translation of MashTag in SunTag frame.
7. Skipping of uORF AUG; skipping of SunTag frame AUG; initiation on MoonTag frame AUG; translation of MashTag in MoonTag frame.
8. Skipping of uORF AUG; skipping of SunTag frame AUG; skipping of MoonTag frame AUG; downstream initiation in blank frame; translation of MashTag in blank frame.
9. Skipping of uORF AUG; skipping of SunTag frame AUG; skipping of MoonTag frame AUG; downstream initiation in MoonTag frame; translation of MashTag in MoonTag frame.
10. Skipping of uORF AUG; skipping of SunTag frame AUG; skipping of MoonTag frame AUG; downstream initiation in SunTag frame; translation of MashTag in SunTag frame.

The following equations were used to calculate the fraction of ribosomes in each path.

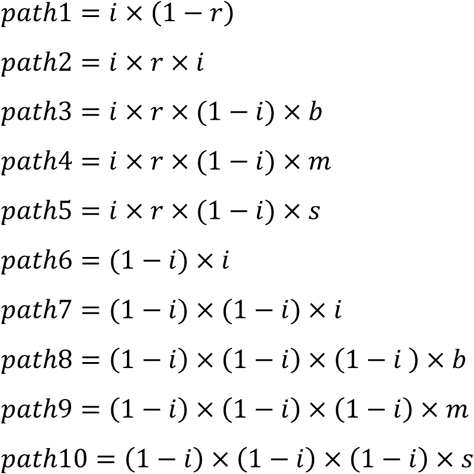

i = probability to initiate translation on an AUG start codon in optimal context.

r = probability to reinitiate scanning after uORF translation.

b, m, or s = probability to initiate translation on a near-cognate start site in either the blank (b), the MoonTag (m), or the SunTag (s) frame downstream of the MoonTag AUG start site

Multiple different paths lead to translation signal of one of the three frames. Therefore:

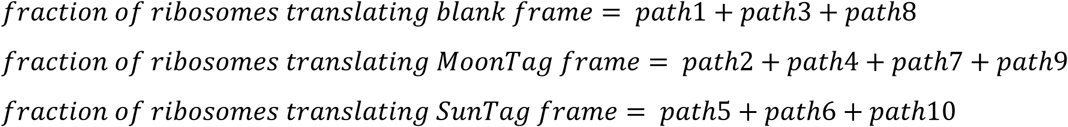

To calculate the values for the different constants (i, r, n) and the fraction of ribosomes in each path, the following assumptions were made:

1. The probability of a scanning ribosome to initiate translation on an AUG in optimal context (i.e. GCCACC<u>AUG</u>G) that is encountered, is constant irrespective of the position of the AUG in the mRNA.
2. The probability to initiate on a near-cognate start site downstream of the MoonTag AUG start site is equal for all three frame (MoonTag, SunTag, and blank frame).
3. The combined translation initiation rate of the MoonTag, SunTag, and blank frame is equal for the MoonStart and uORF reporter.

To determine the probability of a ribosome initiating translation on an AUG start site in optimal context (i), we examined the frequency of translation initiation at the AUG of the MoonStart reporter. The median initiation rate in the MoonTag frame on the MoonStart reporter was 0.88 ribosomes/min, which represents a combination of translation initiation on the MoonTag AUG start codon, and skipping of the AUG and downstream initiation in the MoonTag frame. While we cannot de-convolve these two paths directly, we could measure the translation initiation rate in the SunTag frame (0.1 ribosomes/min), which represents leaky scanning of the MoonTag AUG and downstream initiation on a near-cognate start site in the SunTag frame. One of our assumptions is that the frequency of initiation on near-cognate start sites downstream of the MoonTag AUG is equal in all three frames. Based on this assumption b=m=s=0.33. Thus, if 0.1 ribosomes/min initiate on a near-cognate start site in the SunTag frame, we assume that 0.1 ribosomes/min are also initiating in the MoonTag frame on a near-cognate start site, leaving 0.88-0.1 = 0.78 ribosomes/min for initiation on the MoonTag AUG. Addition of a similar blank frame initiation rate (0.1 ribosomes/min) results in an overall initiation rate of 0.1 + 0.1 + 0.1 + 0.78 = 1.08 for all three frames combined. 0.78 of 1.08 ribosomes/min are initiating on the MoonTag AUG, which represents 0.78 / 1.08 = 0.73 of initiating ribosomes. Therefore i = 0.73, which is comparable to a previously determined probability of initiating on an AUG start codon with optimal context (i = 0.86 based on (Ferreira et al., 2013)).

Finally, we determined the remaining constant r, the probability to reinitiate scanning after uORF translation. To calculate r, we used the following equation:

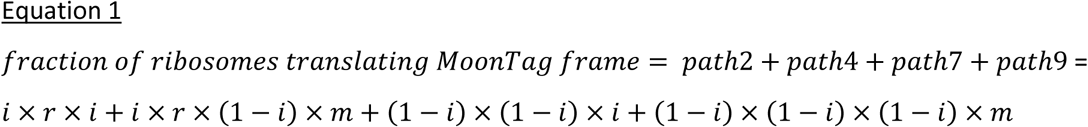

To solve this equation, we first calculated the fraction of ribosomes translating the MoonTag frame. The median MoonTag translation rate on the uORF reporter was 0.36 ribosomes/min. As we assumed that the combined translation initiation rate is equal for the MoonStart and uORF reporter, and we had already determined the total initiation rate for the MoonStart reporter to be 1.08 ribosomes/min, the total translation rate on the uORF reporter is also 1.08 ribosomes/min and thus the relative fraction of ribosomes translating the MoonTag frame is 0.36/1.08 = 0.27. Using this value, we could solve equation 1, which led to r = 0.54.

In parallel, r was calculated based on the total blank frame translation rate, using the following equation:

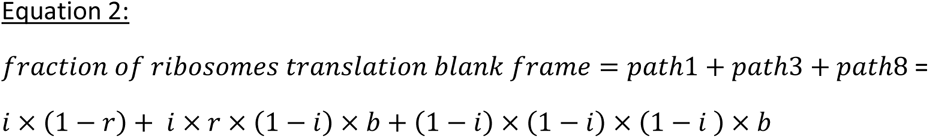

To solve this equation, we first the fraction of ribosomes translating the blank frame, based on the the assumption that combined translation initiation rate is equal for the MoonStart and uORF reporter (1.08 ribosomes/min). The median MoonTag (0.36 ribosomes/min) and SunTag translation (0.29 ribosomes/min) rates were then used to calculate the blank frame translation rate: 1.08-0.36-0.29=0.42 ribosomes/min. The relative fraction of ribosomes translation the blank frame is therefore: 0.42/1.08=0.39. Using this value, we could solve equation 2, and determine r=0.40.

To calculate the fraction of ribosomes per path, the mean of the two values for r that were calculated by solving equations 1 and 2 (0.47) was used.

To confirm these constants, we took advantage of the experiment in which we removed the stop codon of the uORF. Based on the constants, we predicted the effects of removing the uORF stop codon on the translation paths and the translation rates per frame. Removal of the stop codon essentially prevents reinitiation and thus leads to an r = 0 value. With this new value for r, the fraction of ribosomes translating each translation path was computed. Then, the translation rate of the MoonTag and SunTag combined was calculated (r=0; 0.29 ribosomes/min). Similarly, the translation rate of SunTag and MoonTag combined was calculated in the presence of the uORF stop codon (r=0.47; 0.66 ribosomes/min). These calculation revealed that the removal of the uORF stop codon is predicted to induce a 56% (from 0.66 to 0.29 ribosomes/min) reduction in the translation rate. When measuring the translation rates on the uORF reporter in the presence or absence of the uORF stop codon, we observed 58% reduction (from 0.80 to 0.33 ribosomes/min) in median translation rates, validating our calculated parameters.

### 4.5 Statistical analysis

#### 4.5.1 Statistics of population differences

To compare different data sets, students’ T tests or Mann-Whitney tests were performed, as indicated in the figure legends.

##### 4.5.2 Calculating p-values for differences in OOF translation frequencies among different mRNAs

To test whether observed differences in the amount of OOF translation on different MashTag reporter mRNAs could be explained by chance, the OOF translation frequency of individual mRNAs was compared to the OOF translation frequency of the total population of mRNAs. Per mRNA a binomial test was conducted, using the total number of ribosomes on each mRNA, the number of ribosomes translating the SunTag frame on the same mRNA and the median SunTag translation frequency on all mRNAs. Note that this analysis had limited statistical power to detect MoonStart mRNAs with increased MoonTag translation; based on analysis of all mRNAs, 93% of ribosomes initiate translation in the MoonTag frame on the MoonStart reporter. Even if all ribosomes initiate in the MoonTag frame during our observation period (~25 min, ~25-50 initiating ribosomes) the bias towards the MoonTag start site is only moderately statistically significant, so much longer traces are needed to achieve strongly significant p-values for enrichment of MoonTag translation. Nonetheless, we do find 2/85 mRNAs with an enrichment in MoonTag translation initiation events with a p-value > 0.01 and 8/85 mRNAs with a p-value > 0.05.

##### 4.5.3 Linear regression analysis

To perform linear regression analysis on SunTag and MoonTag fluorescence over time, individual intensity traces were first smoothed by generating moving averages of 3 consecutive time points to reduce the noise in intensity traces. For every mRNA, MoonTag intensities were then plotted against SunTag intensities for each time point and linear regression analysis was performed on the SunTag and MoonTag intensity data, and the R^2^-value was determined for each mRNA. Only mRNAs with at least one MoonTag and one SunTag ribosome were included in the analysis.

##### 4.5.4 Calculating sliding window p-values

To test whether changes in the usage of translation initiation sites occurred during the life-time of a mature mRNA, a statistical test was designed, referred to as a sliding window approach (also see Fig. S4A). First, ribosomes were fit to raw intensity traces, and the time of each translation initiation event was determined, as described in section 3.1. Second, initiation events in both MoonTag and SunTag frames were merged onto a single time-line. Third, a ‘window’ of 10 consecutive initiation events was defined as the first 10 initiation events of the trace. Additional windows were defined by sliding the window along all initiation events, moving the window one initiation event per iteration. Note that mRNA traces with 10 or less initiation events were excluded from analysis, as we could not generate multiple windows on these mRNAs. In this way, a collection of windows was created along the trace of an mRNA, in which each window represented 10 consecutive initiating ribosomes. For each window, the number of SunTag initiation events was determined. Fourth, the relative SunTag initiation frequency in each window was compared to the SunTag initiation frequency of the entire mRNA to test whether the window contained an increase or decrease in the number of SunTag initiation events as compared to the total trace. To provide a significance value for each window, a binomial test was performed. Finally, the p-value was determined for each window of an mRNA and the lowest p-value per mRNA was determined.

**Table S1.** number of repeats, cells, and mRNAs per experiment. Overview of experimental repeats and mRNAs analyzed per experiment. (Re)plotting data from same experiment means either the same data is plotted again (as indicated in Figure legend) or same data is used for another type of analysis. Colored box indicates that same data points have been used multiple times and color of box refers to same data set.

| Figure | Description | repeats | cells | mRNAs or events | (re)plotting data from same experiment |
| --- | --- | --- | --- | --- | --- |
| 1D | 4xMoonTag-11aalinker | 1 | 32 | 32 |  |
|  | 12xMoonTag-11aalinker | 1 | 26 | 26 |  |
|  | 24xMoonTag-11aalinker | 1 | 23 | 23 |  |
|  | 24xMoonTag-5aalinker | 1 | 25 | 25 |  |
| 1I | 24xMoonTag-kif18b-STOP-24xSunTag-BFP | 4 | 16 | 74 |  |
| 2H | 5'UTR_Emi-Sstart-36xMash1-BFP<br>-3xStop-24xPP7 | 4 | 8 | 17 | * |
| 2I | Mstart-36xMash1-BFP-3xStop-24xPP7 | 3 | 10 | 17 | * |
| 3C | 24xSunTag-kif18b-stop-24xPP7 | 2 | 12 | 54 |  |
|  | 24xMoonTag-kif18b-stop-24xPP7 | 2 | 14 | 61 |  |
|  | 12xMoonTag-12xSunTag-kif18b<br>-stop-24xPP7 | 2 | 8 | 28 |  |
| 4A | Mstart-36xMash1-BFP-3xStop-24xPP7 | 4 | 17 | 85 |  |
| 4C | Mstart-36xMash1-BFP-3xStop-24xPP7 | 4 | 17 | 85 |  |
| 4D | Mstart-36xMash1-BFP-3xStop-24xPP7 | 4 | 17 | 85 |  |
| 4E | 12xMoonTag-12xSunTag-kif18b<br>-stop-24xPP7 | 2 | 8 | 28 |  |
|  | Mstart-36xMash1-BFP-3xStop-24xPP7 | 4 | 17 | 56 | ** |
| 4F | Mstart-36xMash1-BFP-3xStop-24xPP7 | 4 | 17 | 63 | *** |
| 5B | Mstart-36xMash1-BFP-3xStop-24xPP7 | 4 | 17 | 85 |  |
|  | Mstart-Bstart-36xMash1-BFP<br>-3xStop-24xPP7 | 3 | 18 | 71 |  |
|  | Mstart-Mstart-36xMash1-BFP<br>-3xStop-24xPP7 | 2 | 17 | 53 |  |
|  | Mstart-Sstart-36xMash1-BFP<br>-3xStop-24xPP7 | 4 | 16 | 94 |  |
| 5C | Mstart-36xMash1-BFP-3xStop-24xPP7 | 4 | 17 | 85 |  |
|  | ΔSunStop-Mstart-36xMash1<br>-BFP-3xStop-24xPP7 | 4 | 13 | 62 |  |
| 5D | Mstart-36xMash1-BFP-3xStop-24xPP7 | 4 | 17 | 85 |  |
|  | HMBS_5'UTR-Mstart-36xMash1-BFP-3xStop-24xPP7 | 3 | 15 | 54 |  |
|  | RPL12_5'UTR-Mstart-36xMash1-BFP-3xStop-24xPP7 | 3 | 13 | 53 |  |
| 6E | uORF(Blank)-Sstart(in_uORF)-Mstart(reint)<br>-36xMash-BFP-3xstop-24xPP7 | 2 | 10 | 53 |  |
| 6F | 12xMoonTag-12xSunTag-kif18b-stop-24xPP7 | 2 | 8 | 28 |  |
|  | uORF(Blank)-Sstart(in_uORF)-Mstart(reint)<br>-36xMash-BFP-3xstop-24xPP7 | 2 | 10 | 44 | ** |
| 6G | uORF(Blank)-Sstart(in_uORF)-Mstart(reint)<br>-36xMash-BFP-3xstop-24xPP7 | 2 | 10 | 37 | *** |
| S1D | 24xMT-kif18b-STOP-24xST-BFP | 4 | 16 | 74 |  |
| S2A | Sstart-36xMash1-BFP-3xstop-24xPP7<br>+Harr | 3 | 8 | 26 |  |
| S2B | Mstart-24xMash1-BFP-CAAX(Sun) | 2 | 59 | 505 |  |
| S2C | Sstart-36xMash1-BFP-CAAX(Sun) | 2 | 34 | 407 |  |
|  | Mstart-36xMash1-BFP-CAAX(Sun) | 2 | 33 | 399 |  |
| S3A | Mstart-36xMash1-BFP-3xStop-24xPP7 | 4 | 17 | 85 |  |
| S3B | Sstart-36xMash1-BFP-3xstop-24xPP7 | 4 | 17 | 43 |  |
| S3C | Mstart-36xMash1-BFP-3xStop-24xPP7 | 5 | 36 | 195 |  |
| S5A | ΔSunStop-Mstart-36xMash1<br>-BFP-3xStop-24xPP7 | 4 | 13 | 62 |  |
|  | pSB264+DMSO | 4 | 13 | 58 |  |
|  | pSB264+RocA | 3 | 14 | 51 |  |
| S5B | ΔSunStop-Mstart-36xMash1<br>-BFP-3xStop-24xPP7 | 4 | 13 | 62 |  |
|  | pSB264+DMSO | 4 | 13 | 58 |  |
|  | pSB264+RocA | 3 | 14 | 51 |  |
| S5C | Mstart-36xMash1-BFP-3xStop-24xPP7 | 4 | 17 | 85 |  |
|  | HMBS_5'UTR-Mstart-36xMash1-BFP-3xStop-24xPP7 | 3 | 15 | 54 |  |
|  | RPL12_5'UTR-Mstart-36xMash1-BFP-3xStop-24xPP7 | 3 | 13 | 53 |  |
| S6A | uORF(Blank)-Sstart(in_uORF)-Mstart(reint)-36xMash-BFP-3xstop-24xPP7 | 2 | 10 | 53 |  |
|  | uORF(Blank)-Mstart(in_uORF)-Sstart(reint)-36xMash-BFP-3xstop-24xPP7 | 3 | 9 | 34 |  |
| S6B | Mstart-36xMash1-BFP-3xStop-24xPP7 | 4 | 17 | 85 |  |
|  | uORF(Blank)-Sstart(in_uORF)-Mstart(reint)-36xMash-BFP-3xstop-24xPP7 | 2 | 10 | 53 |  |
|  | uORF(Blank)_nouORFStop-Sstart(in_uORF)-Mstart(reint)-36xMash-BFP-3xstop-24xPP7 | 3 | 17 | 45 |  |
| S6C | Mstart-36xMash1-BFP-3xStop-24xPP7 | 4 | 17 | 85 |  |
|  | uORF(Blank)-Sstart(in_uORF)-Mstart(reint)-36xMash-BFP-3xstop-24xPP7 | 2 | 10 | 53 |  |
|  | uORF(Blank)_nouORFStop-Sstart(in_uORF)-Mstart(reint)-36xMash-BFP-3xstop-24xPP7 | 3 | 17 | 45 |  |
| S6D | Mstart-36xMash1-BFP-3xStop-24xPP7 | 4 | 17 | 85 |  |
|  | uORF(Blank)-Sstart(in_uORF)-Mstart(reint)-36xMash-BFP-3xstop-24xPP7 | 2 | 10 | 53 |  |
|  | uORF(Blank)_nouORFStop-Sstart(in_uORF)-Mstart(reint)-36xMash-BFP-3xstop-24xPP7 | 3 | 17 | 45 |  |

\*:only included OOF events that fullfill criteria (see STAR Methods)
\*\*: only included mRNAs with initiation events in both reading frames
\*\*\*:only used mRNAs with >10 initiation events during timelaps

**Table S2.**
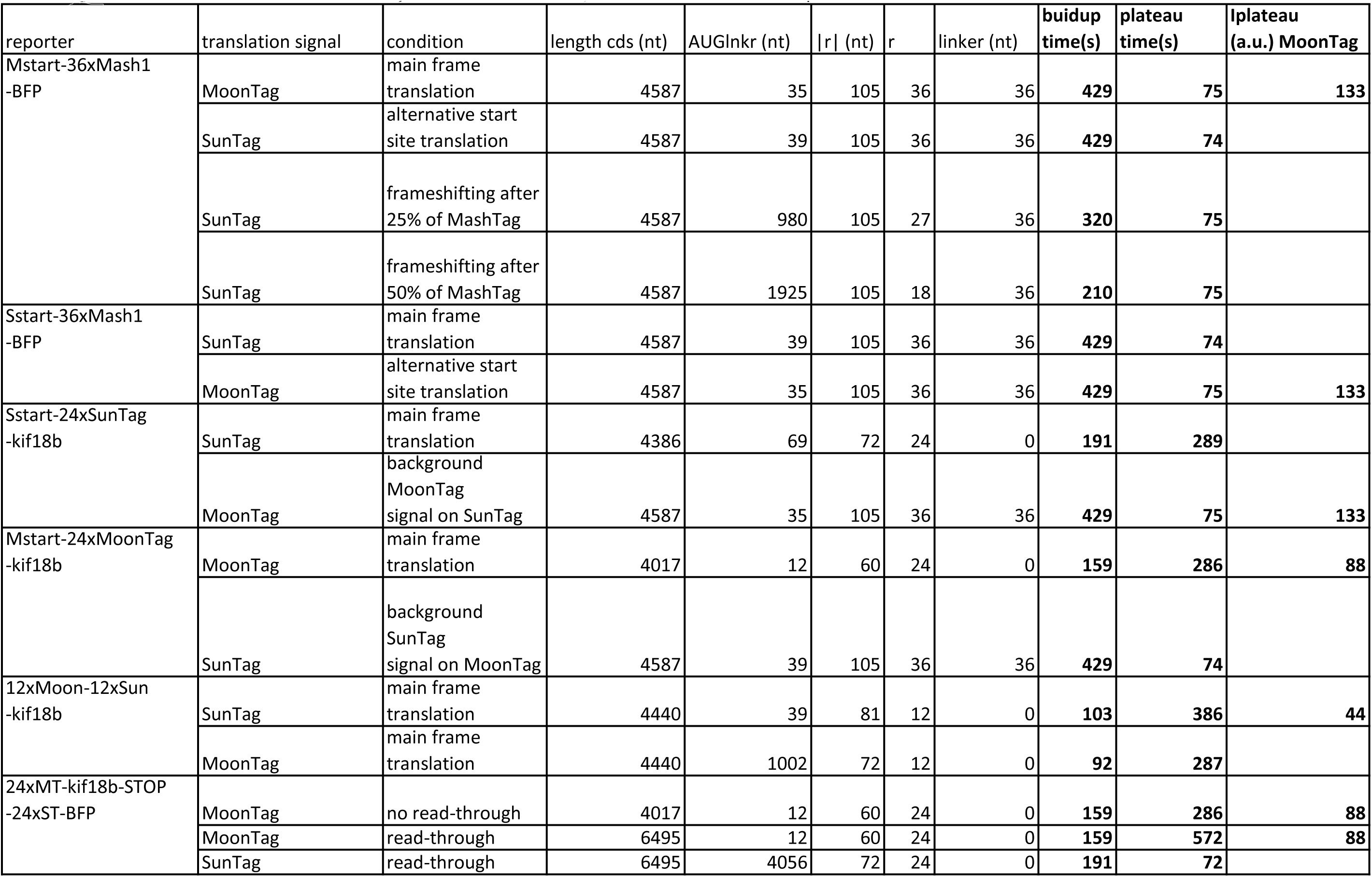
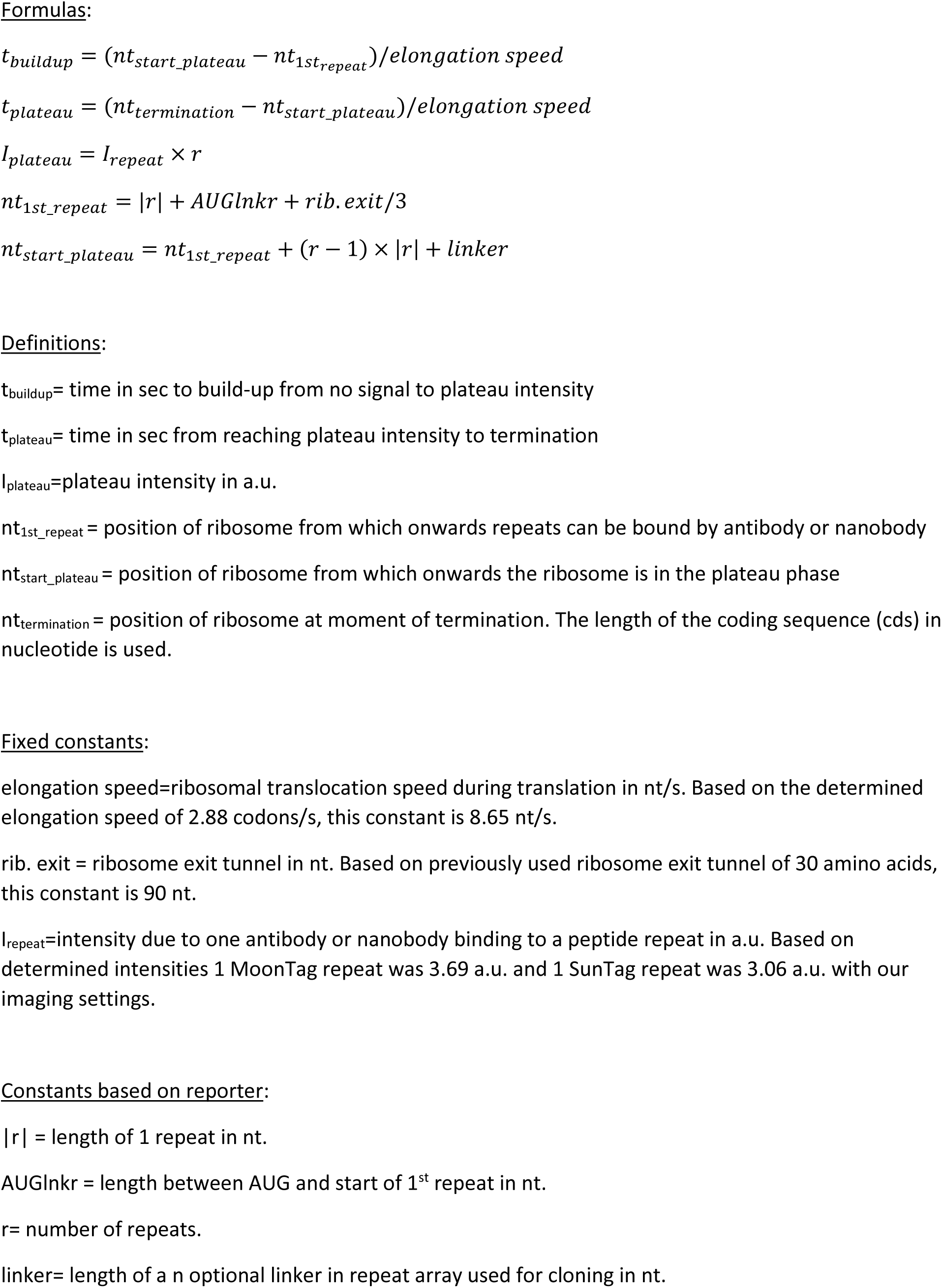
Translation parameters used to define theoretical single ribosome traces per reporter. Based on the parameters per reporter, and the displayed formulas we calculated the build-up time, plateau time, and plateau intensity (bold) of a single ribosome. For each reporter, the theoretical single ribosome values were calculated for both the MoonTag and the SunTag signals. Note that on the 24xMoonTag reporter, the SunTag single ribosome trace of 36xMashTag was used as SunTag signal. Similarly, the MoonTag single ribosomes trace of 36xMashTag was used as MoonTag signal on the 24xSunTag reporter. For details see Methods section 3.1 and 3.3.

**Movie S1 - related to figure 1. Real-time observation of 3’UTR translation of a single mRNA molecule.**

A Moon/Sun cell expressing the read-through reporter (indicated in Fig. 1H). Images were acquired every 30 s (movie duration is 6.5 min) on a spinning disk confocal microscope focusing near the bottom plasma membrane of the cell. One of the many MoonTag translating mRNAs (blue) shows SunTag signal (green). Movie field of view is 4.6 × 4.6 µm.

**Movie S2 - related to figure 2. Real-time observation of OOF translation on a single MashTag reporter mRNA.**

A Moon/Sun cell expressing the MashTag reporter (indicated in Fig. 2B). Images were acquired every 30 s (movie duration is 9 min) on a spinning disk confocal microscope focusing near the bottom plasma membrane of the cell. One of the two MoonTag translating mRNAs (blue, lower mRNA) also shows SunTag signal (green), indicating OOF translation. Movie field of view is 5.4 × 5.4 µm.

**Figure S1 – Related to Figure 1.**
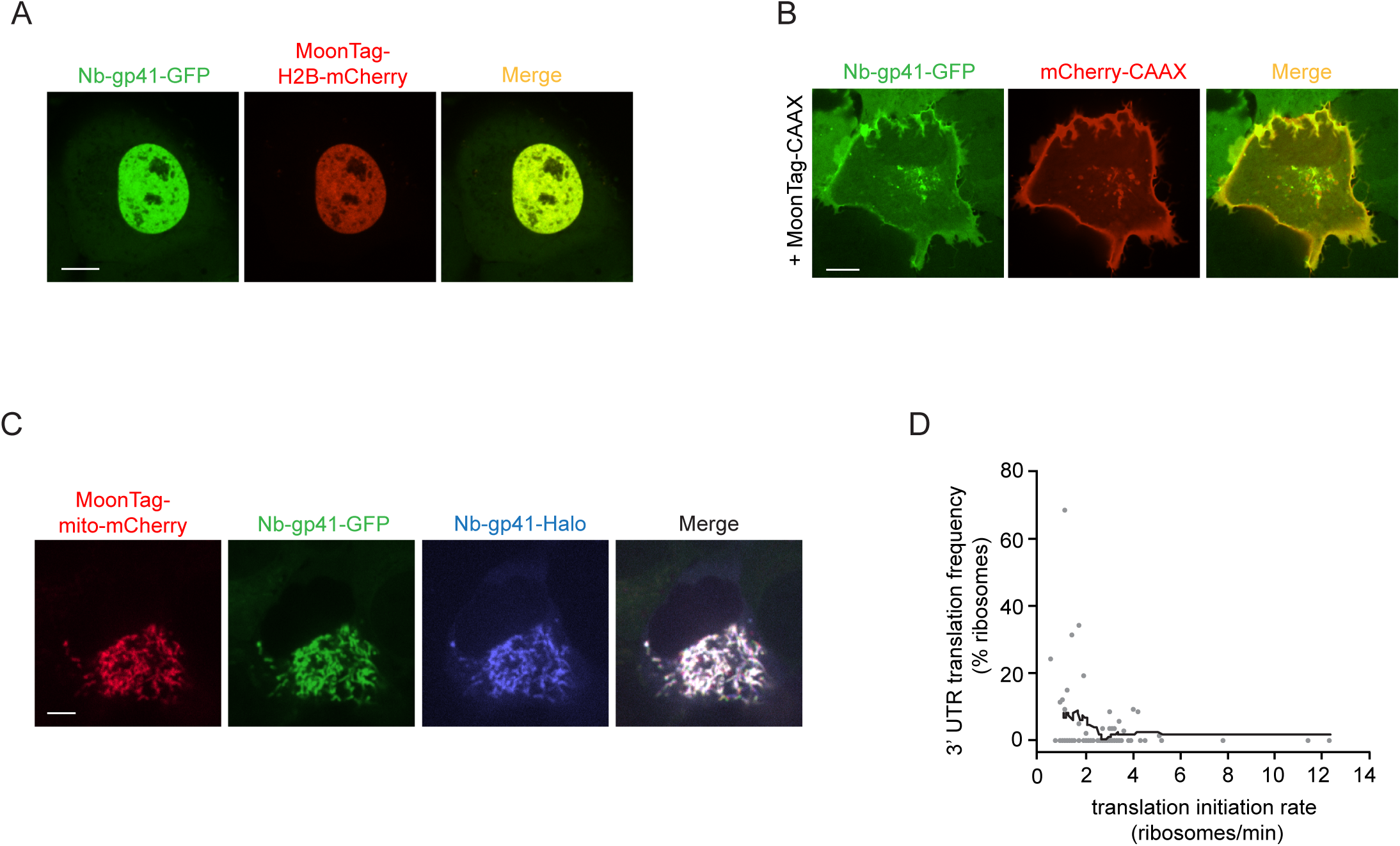
A-C) U2OS cells stably expressing MoonTag nanobody-GFP were transfected with 12xMoonTag-H2B-mCherry (A), 12xMoonTag-CAAX and mCherry-CAAX (B), or 12xMoonTag-Mito-mCherry and MoonTag nanobody-Halo^JF646^ (C). Representative cells are shown. Scale bars, 10µm (A, B), and 5µm (C). D) Moon/Sun cells expressing the reporter indicated in Fig. 1H. Correlation between the coding sequence translation initiation rate and 3’UTR translation frequency on single mRNAs is shown. Every dot represents a single mRNA and line depicts moving average over 15 mRNAs (See Methods).

**Figure S2 – Related to Figure 3.**
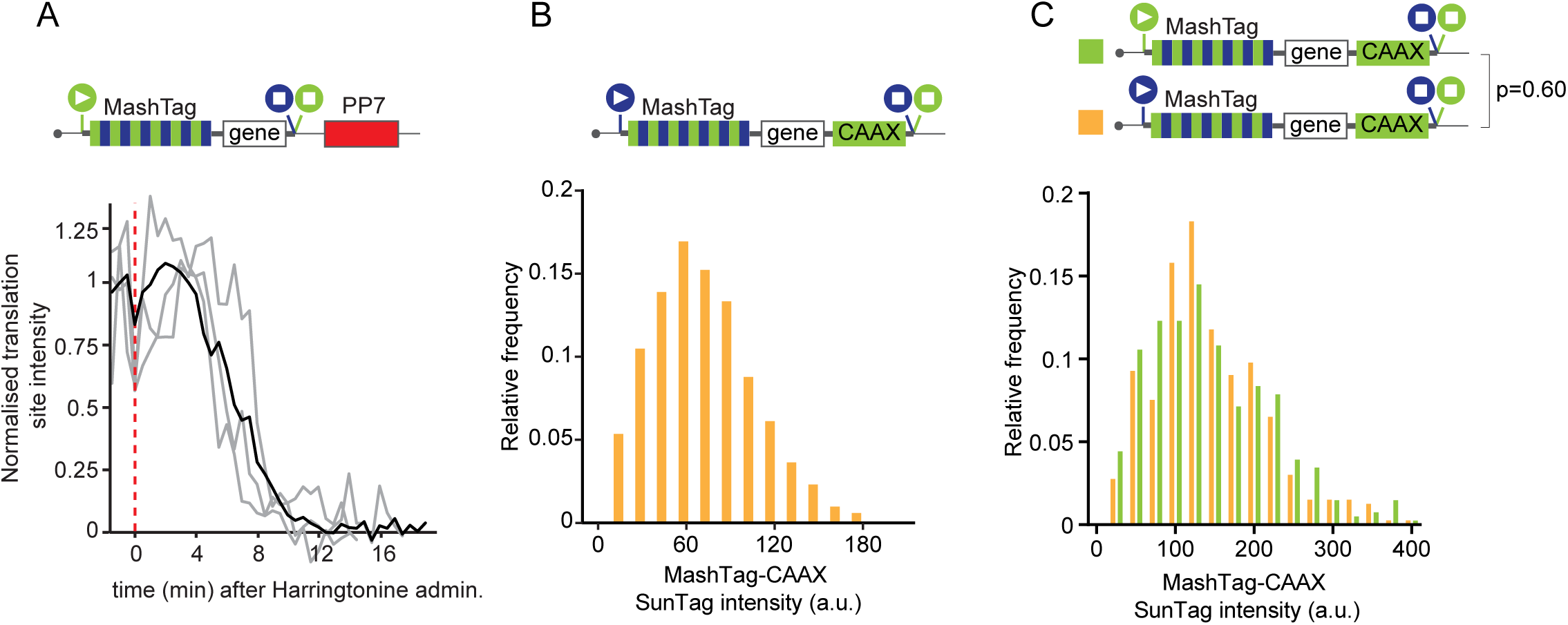
A-C) Schematics: filled circles with white triangles represent translation start sites; filled circles with white squares represent translation stop codons. Colors of the filled circles indicate reading frame; blue indicates MoonTag, while green represents SunTag reading frame. Indicated reporters were transfected into Moon/Sun cells. A) Normalized SunTag intensity on mRNAs after harringtonine treatment. Grey lines depict single example mRNA traces and the black line shows the average of all mRNAs. Red line indicates harringtonine addition. B) Distribution of the intensity of mature proteins expressed from the SunTag frame. Mature proteins were tethered to the plasma membrane through a CAAX motif. C) Distribution of the intensity of mature proteins expressed in the SunTag frame. SunTag proteins were expressed either from the main reading frame (green) or as OOF translation protein products (orange). Mature proteins are tethered to the membrane through a CAAX domain encoded in the SunTag frame. P-value is based on a two-tailed Student’s T-Test.

**Figure S3 – Related to Figure 3.**
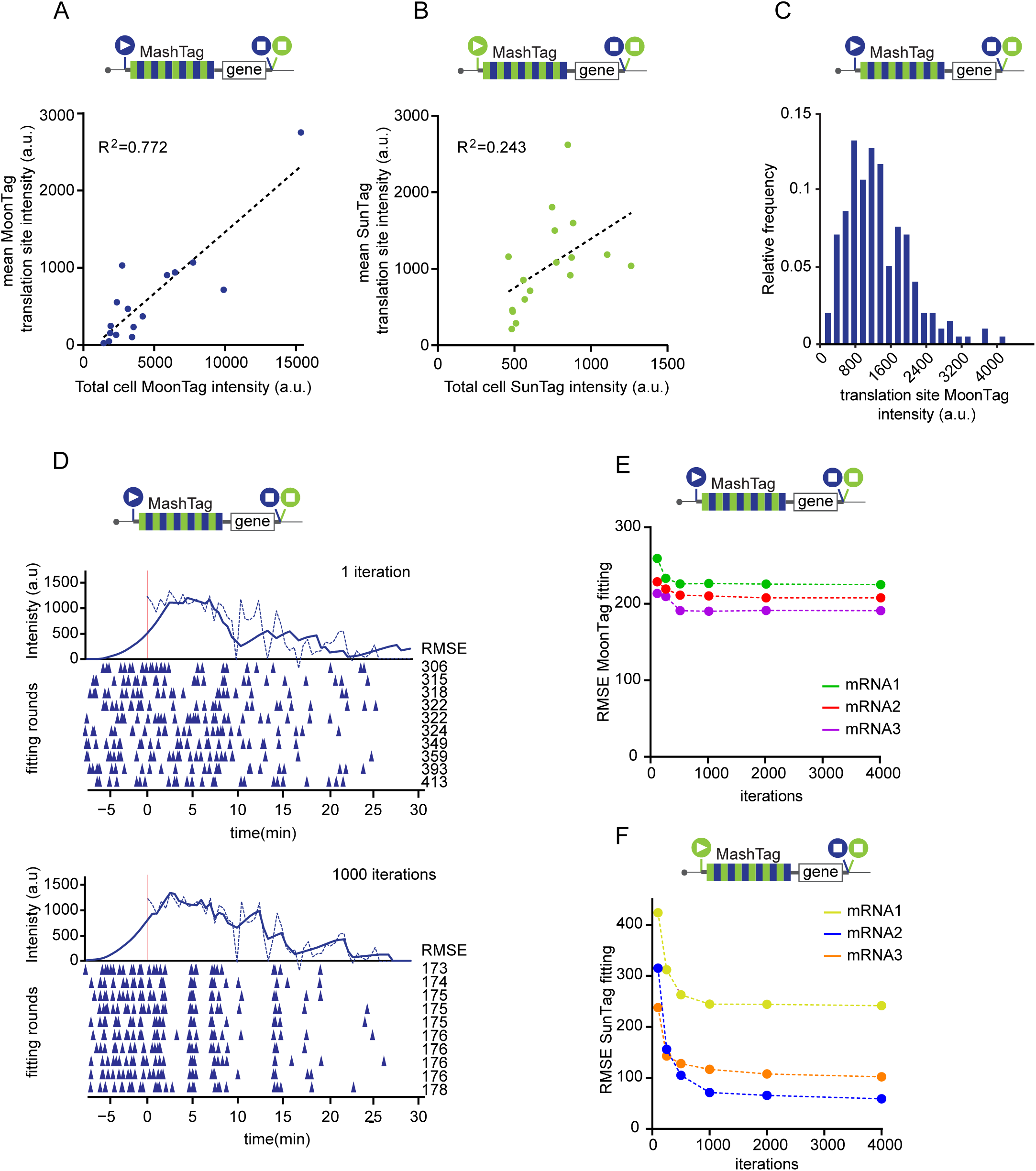
A-F) Schematics: filled circles with white triangles represent translation start sites; filled circles with white squares represent translation stop codons. Colors of the filled circles indicate reading frame; blue indicates MoonTag, while green represents SunTag reading frame. For simplicity, 24xPP7 sites in the 3’UTR are not depicted in the schematics. Moon/Sun cells were transfected with indicated reporters. A) Correlation between total cell MoonTag intensity and average MoonTag translation signal on mRNAs. B) Correlation between total cell SunTag intensity and average SunTag translation signal on mRNAs. C) Distribution of the intensity of the MoonTag translation signal of individual mRNAs. D) An example intensity track (dashed line) and fit (solid line) are shown for the MoonTag signal after one iteration (top) or 1000 iterations (bottom) of fit optimization. Colored triangles below the x-axes represent translation initiation events. Each row of triangles illustrates an independent round of fitting. Corresponding root mean squared error (RMSE) values for each round of fitting are shown. (E-F) RMSEs after indicated number of iterations of fit optimization is shown for three representative mRNAs.

**Figure S4 – Related to Figure 4.**
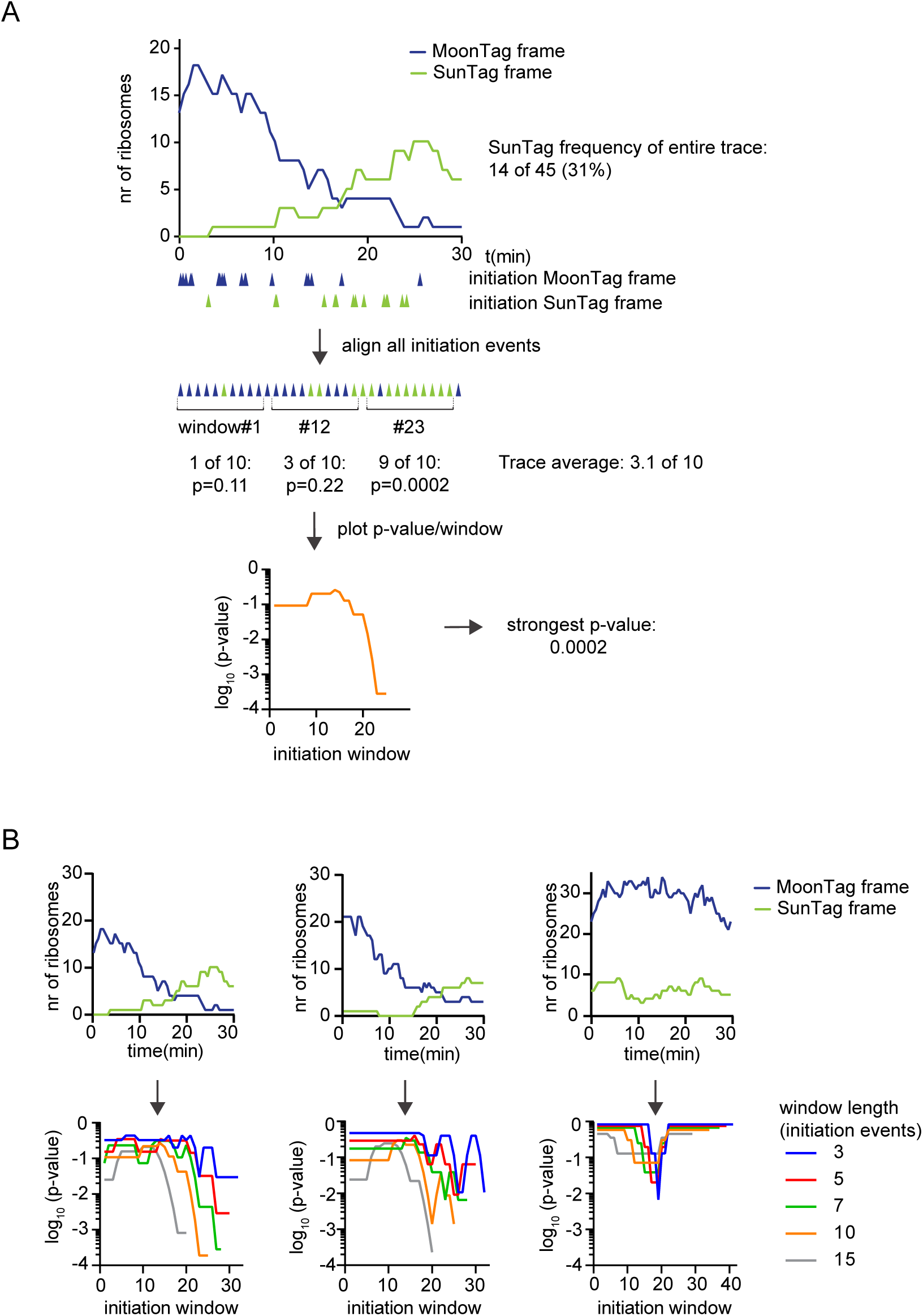
Moon/Sun cells expressing the reporter indicated in Fig. 4A were analyzed. A) Schematic of sliding window analysis approach. First, ribosomes were fit to raw intensity traces, and the time of each translation initiation event was determined (indicated by triangles under the x-axis), as described in Fig. 3B (top). Initiation events in both MoonTag and SunTag frames were then merged onto a single time-line. For each consecutive set of initiation events (window length of 10 initiation events is shown), a p-value was calculated (binomial test), which represents the probability of observing the ratio between MoonTag and SunTag initiation events within that window, based on the MoonTag and SunTag translation initiation rates of the entire mRNA trace (middle). The p-value for each window of 10 consecutive initiation events of an mRNA was plotted and the strongest p-value per mRNA was determined (bottom). B) Example graphs of the number of ribosomes in each reading frame over time for 3 mRNAs (top panel). Corresponding sliding window p-value plots are shown below for sliding windows with indicated number of initiation events per window. The mRNA displayed here was also used as an example in Fig. 4F.

**Figure S5 – Related to Figure 5.**
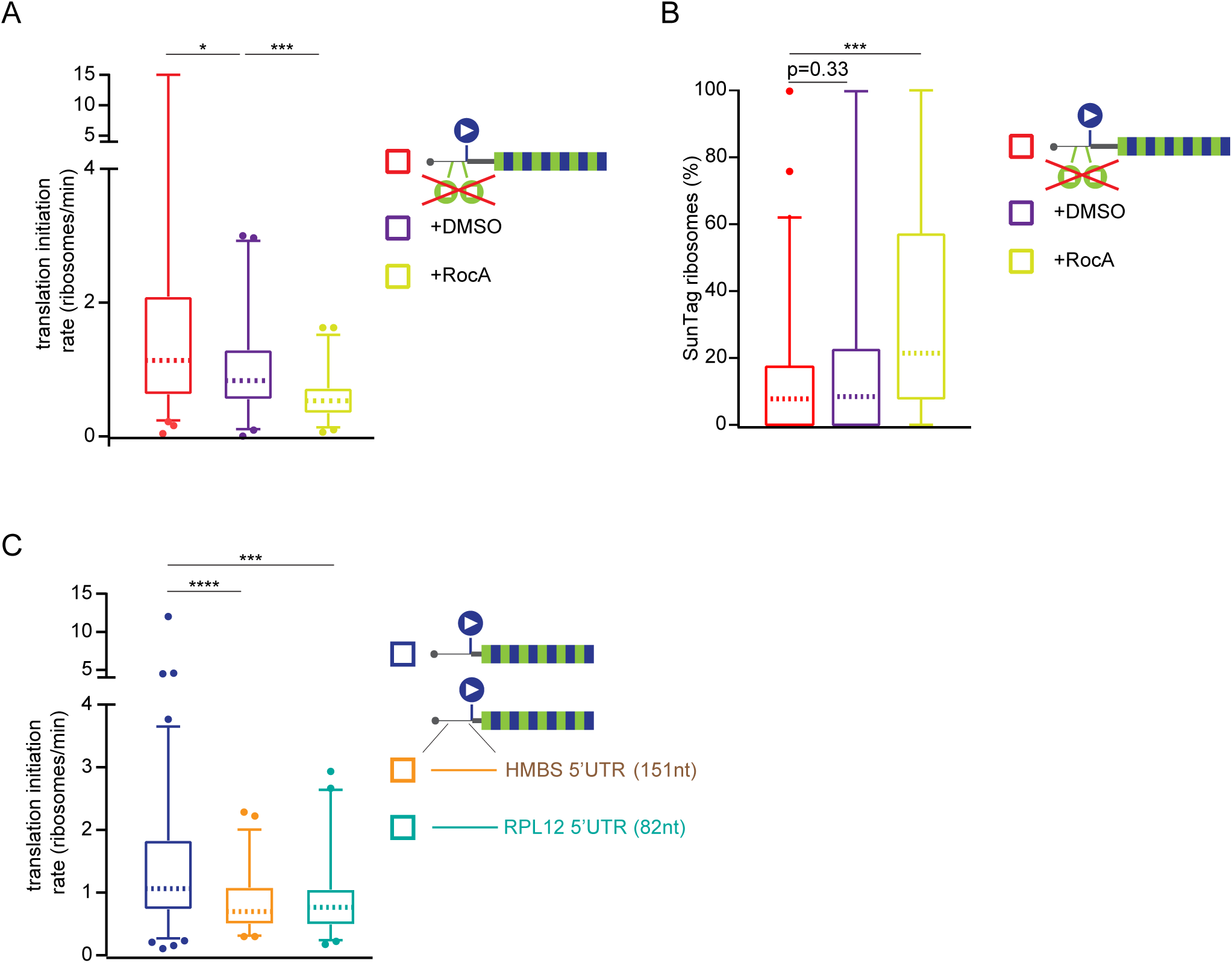
A-C) Schematic reporter: filled circles with white triangles represent translation start sites; filled circles with white squares represent translation stop codons. Colors of the filled circles indicate reading frame; blue indicates MoonTag, green indicates SunTag reading frame. For simplicity, reporter schematics only show 5’ region of the mRNA. Indicated reporter mRNAs were expressed in Moon/Sun cells. Boxplots are shown of the overall translation initiation rates (i.e. initiation rates of MoonTag and SunTag frames combined) (A, C) or fraction of SunTag translation (B) for single mRNAs. P-values are based on two-tailed Mann-Whitney tests; * p<0.05; ** p<0.01; *** p<0.001; **** p<0.0001. Dashed line represents median value, box indicates 25-75% range, and whiskers indicated 5-95% range. Number of experimental repeats and mRNAs analyzed per experiment are listed in Table S1.

**Figure S6 – Related to Figure 6.**
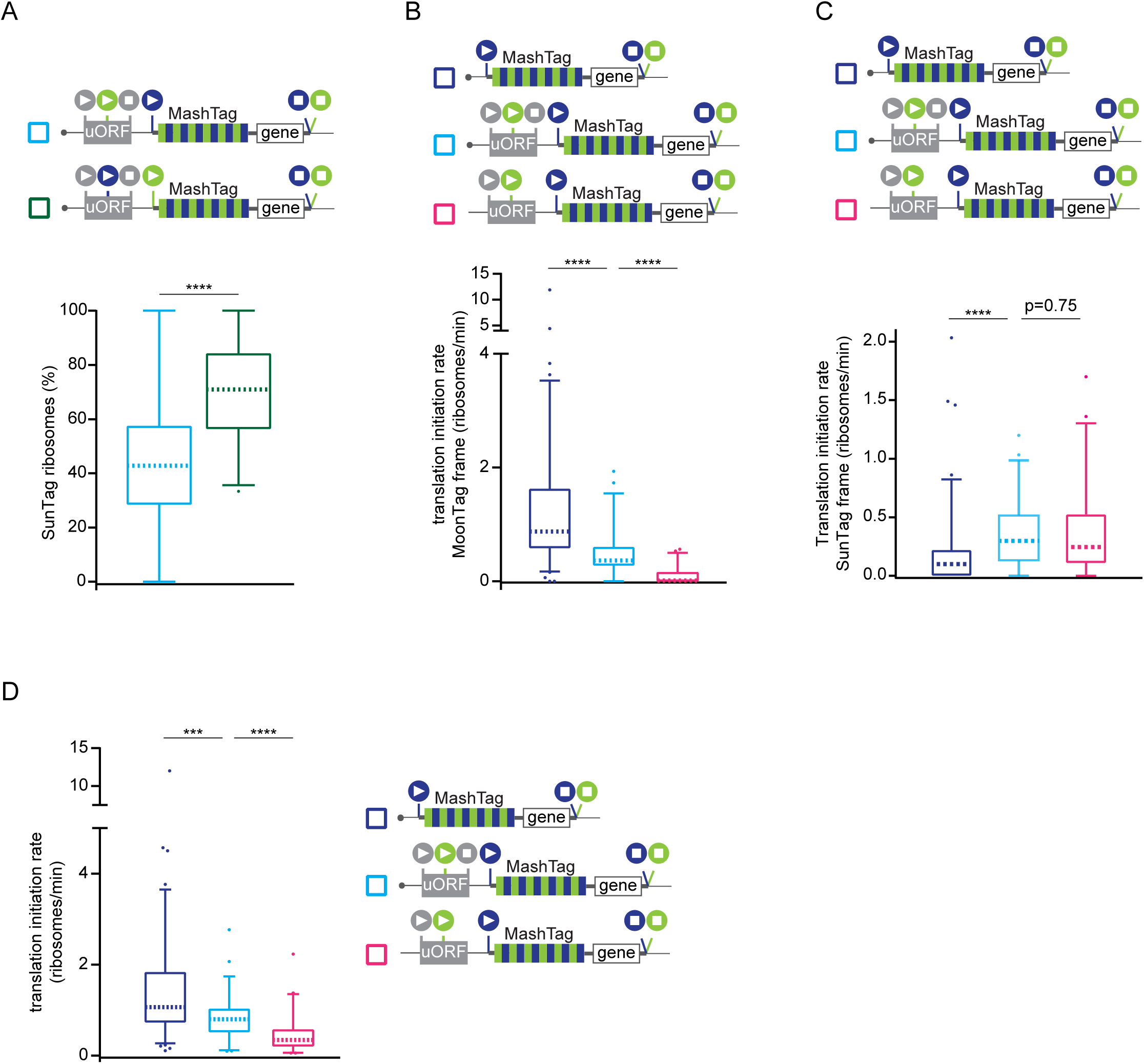
A-D) Schematic reporter: filled circles with white triangles represent translation start sites; filled circles with white squares represent translation stop codons. Colors of the filled circles indicate reading frame; blue indicates MoonTag, green indicates SunTag, and grey indicates blank reading frame. For simplicity, 24xPP7 sites in the 3’UTR are not depicted in the schematics. Indicated reporter mRNAs were expressed in Moon/Sun cells. A) Boxplots of relative initiation frequency in the SunTag frame (relative to the sum of the MoonTag and SunTag frame) on single mRNAs. P-values are based on two-tailed Mann-Whitney tests: *** p<0.001; **** p<0.0001. B-D) Boxplots of translation initiation rates in the MoonTag frame (B), SunTag frame (C), or MoonTag and SunTag frame combined (D) for single mRNAs. Dashed line represents median value, box indicates 25-75% range, and whiskers indicated 5-95% range. Number of experimental repeats and mRNAs analyzed per experiment are listed in Table S1.

